# Evolution of a pest: towards the complete neuroethology of *Drosophila suzukii* and the subgenus *Sophophora*

**DOI:** 10.1101/717322

**Authors:** Ian W. Keesey, Jin Zhang, Ana Depetris-Chauvin, George F. Obiero, Markus Knaden, Bill S. Hansson

**Affiliations:** Max Planck Institute for Chemical Ecology, Department of Evolutionary Neuroethology, Hans-Knöll-Straße 8, D-07745 Jena, Germany; Department of Biochemistry and Biotechnology, Technical University of Kenya, Haille-Sellasie Avenue, Workshop Rd, 0200, Nairobi, Kenya

## Abstract

Comparative analysis of multiple genomes has been used extensively to examine the evolution of chemosensory receptors across the genus *Drosophila*. However, few studies have delved into functional characteristics, as most have relied exclusively on genomic data alone, especially for non-model species. In order to increase our understanding of olfactory evolution, we have generated a comprehensive assessment of the olfactory functions associated with the antenna and palps for *Drosophila suzukii* as well as several other members of the subgenus *Sophophora*, thus creating a functional olfactory landscape across a total of 20 species. Here we identify and describe several common elements of evolution, including consistent changes in ligand spectra as well as relative receptor abundance, which appear heavily correlated with the known phylogeny. We also combine our functional ligand data with protein orthologue alignments to provide a high-throughput evolutionary assessment and predictive model, where we begin to examine the underlying mechanisms of evolutionary changes utilizing both genetics and odorant binding affinities. In addition, we document that only a few receptors frequently vary between species, and we evaluate the justifications for evolution to reoccur repeatedly within only this small subset of available olfactory sensory neurons.

## INTRODUCTION

One of the advantages of working within the *Drosophila* genus includes the wide-array of evolutionary specializations that these species display, especially in regard to host preference, but also habitat choice, morphology and mate selection^1,2^. As an additional benefit, the genus affords a vast amount of genomic data, which has been generated and accumulated since the early 21^st^ century^3,4^. The last ten years have also brought attention towards another novelty from this group, an agricultural pest species in the form of *Drosophila suzukii*, which has now invaded North and South America, Europe as well as Asia^5–9^. Furthermore, this pest insect has prompted both integrated pest management efforts as well as evolutionary neuroethology research, where the distinct ecological niche of attacking fresh or ripening fruit has afforded a unique opportunity for comparison among the other model species within this well-studied genus of flies, which usually target softened, fermented host resources^10–13^.

Previous research has sought to outline either the olfactory or gustatory systems of several species of *Drosophila*, either for evolutionary or ecological comparisons of host plant, mate preference or reproductive isolation, including *D. melanogaster, D. sechellia*, and *D. simulans*^14–17^. However, only a few studies have conducted electrophysiological assessments of members outside of this subgroup^18^, with most research examining non-*melanogaster* species only in regards to a single, specific olfactory sensory neuron (OSN) of interest, and usually only as a means to show the conserved nature of that functional neuronal type^19–23^. Some more distantly related *Drosophila* species have begun to be examined in more depth, such as *D. mojavensis*^24^, which is a model for incipient speciation; however, this species is a member of an entirely different subgenus, and as such, is quite far removed from the more robust datasets afforded by the *melanogaster* clade. These factors may also mean that these distantly related species are much more difficult to utilize to assess patterns and mechanisms of evolutionary selective pressure, at least in their direct comparison to the established molecular models. Therefore, in contrast, the detailed examination of the non-*melanogaster* members of the *Sophophora* subgenus, which itself includes substantial variation in host and habitat choice, perhaps represents a closer group of relatives to optimize the comparisons of evolutionary variation in chemosensation. Here again, *D. suzukii* and its subgroup offer an ideal model for unraveling the complexities of olfactory evolution, especially given their relative phylogenetic proximity to the principal scientific models within the *melanogaster* clade^1,2^.

A fundamental aspect of studying evolution revolves around understanding the processes by which genetic variation can be generated naturally, such as through mutation, genetic drift, or recombination, and then how selective forces, including environmental, ecological and developmental factors, subsequently act upon this variance to drive speciation and specialization. For many *Drosophila* species, there already exists a robust library of molecular resources, as nearly complete genomic datasets are publically available; however, far less information is available describing the functional components of host ecology, such as olfactory, gustatory, auditory and visual preferences, or describing behavioral or habitat variations between these species. Thus for most members across this incredibly well studied genus, more is known about their genome than their ecology. In an effort to expand our knowledge about these species, here we provide a functional olfactory landscape encompassing 20 different species within the *Sophophora* subgenus, with particular focus on *D. suzukii* and its closest relatives, in order to examine the common variables that are associated with the evolutionary emergence of this insect pest. In addition, we address how protein sequence coding of olfactory receptors correlate with functional evolution of odorant binding and ligand selectivity, here through high-throughput comparison of chemosensory variation overlaid with tertiary protein structures from the available model species. Thus, in summary, the present study establishes the missing ingredients for us to start understanding the mechanisms of olfactory evolution, while also providing critical chemosensory evaluations across 20 species within this highly influential genus of insects.

## RESULTS

### Complete screening of *Drosophila suzukii* olfaction

In order to assess olfactory ligand spectra for all sensillum types, we first revisited the classical model, *D. melanogaster*, where the responses of most sensilla have been previously established^14,24–28^. Here we again mapped out the response profile from each known receptor across the adult fly antenna and palps using a panel of 80 odorants (**Figure 1**). We found that out of 37 unique olfactory sensory neurons (OSNs), only 3 are still currently without a strong ligand candidate (i.e. Or23a, Or2a, Or65a/b/c), although in addition, several co-expressed receptors also have poorly defined response profiles (e.g. Or33a which is found in ab4B). Next, we shifted our focus towards the antenna and palps of *D. suzukii* using the same diverse panel of 80 odorants in order to examine any changes in functional ligand spectra between these two species. Here we determined that of the 37 OSNs found in *D. suzukii*, approximately 86% are functionally conserved between the two species (**Figure 1A**). Moreover, we identified only five OSNs that showed robust functional deviation from the *D. melanogaster* model, including the ab2B-like, ab3A-like, ab9B-like, ab10A-like, and ai3A-like sensory neurons (**Figure 1A**). We also document minor shifts across ab5 and ab7, though not as strong, and we note that we found very few ab8 sensilla across *D.suzukii* trials. Two of the larger deviations in olfactory response for *D. suzukii* have been previously described, including ab2B and ab3A^10^. Interestingly, in the present study we identify two separately responding ab3A OSN types, with an approximately 60:40 split ratio for OSN abundance, where the first (type i) is tuned towards isobutyl acetate (IBA) and the second (type ii) responds most strongly to beta-cyclocitral (βCC). Both of these ab3A type OSNs in *D. suzukii* are tuned quite differently than those found within the corresponding sensillum of *D. melanogaster*, which is tuned instead towards ethyl hexanoate (EH). However, both species (and both type i and type ii, in *D. suzukii*) share identical ligand spectra for the second OSN in the same sensillum ab3B, which responds characteristically towards 2-heptanol as the best ligand. We also found that although the ab2B OSN from *D. suzukii* deviates from *D. melanogaster*, that the ab2A OSN appears identical. As such, we believe these two sensilla in *D. suzukii* (ab2-like and ab3-like) provide the strongest match for the comparative sensilla in D*. melanogaster*, even given the deviations in ligand spectra for select neurons. This idea is further supported by the morphological structure of the sensillum (i.e. large basiconic), and by the large amplitudes that are characteristic of these sensillum types relative to the small basiconics. All these factors combine to strongly-support the identity of these two sensillum types in *D. suzukii* adults, despite the variation in olfactory ligand spectra, as has been previously proposed^10^.

**Figure 1.**
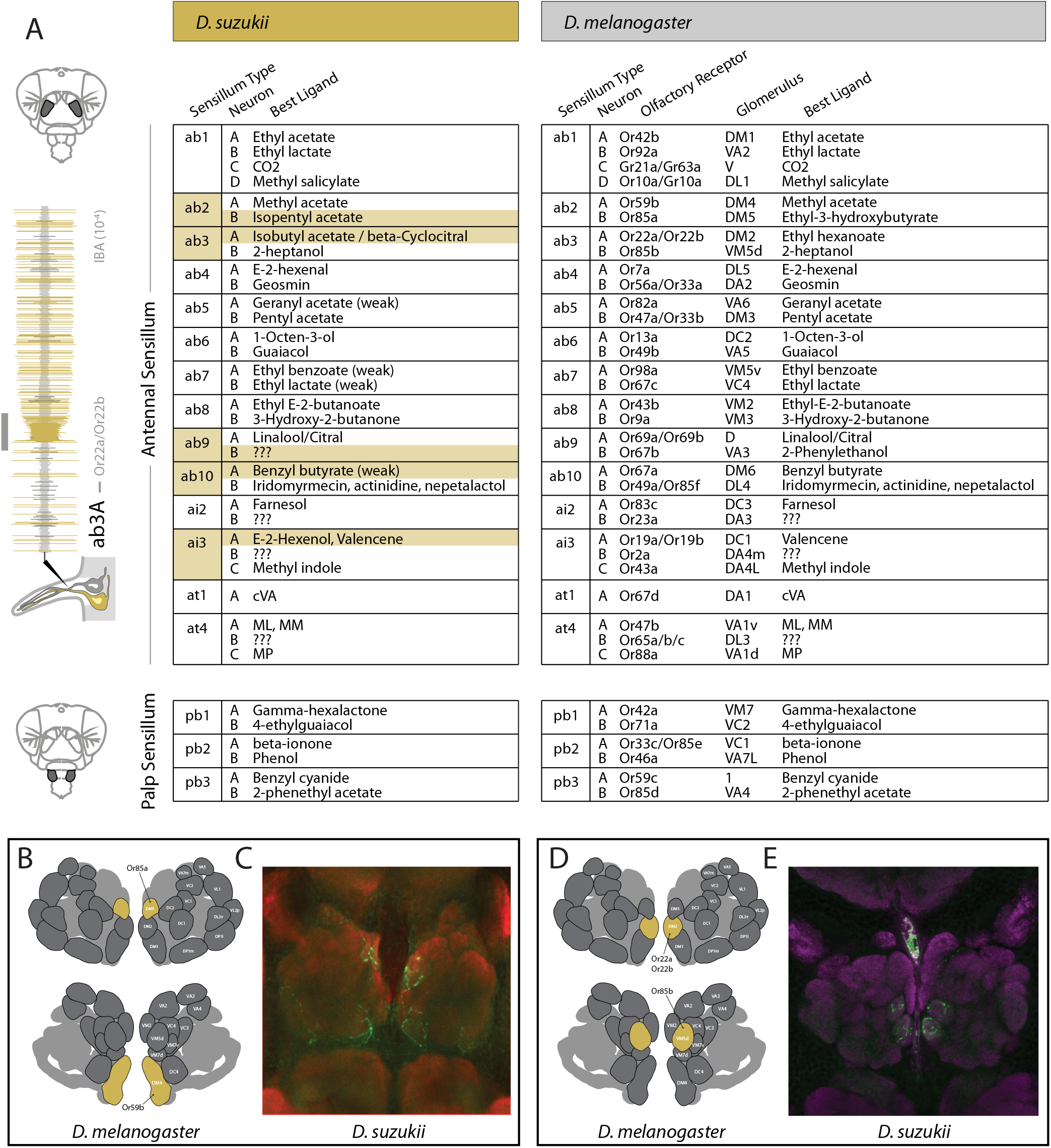
Complete olfactory sensory neuron screen of *Drosophila suzukii*. (A) Shown are the sensillum subtypes, olfactory sensory neurons (OSNs), and best ligands for each of those olfactory channels found within either the antennae or the palps of these two adult *Drosophila* species. For *Drosophila melanogaster*, additional information is given regarding the known olfactory receptor and the corresponding glomerulus for the antennal lobe (AL) for each OSN. In total, 37 neurons were analyzed from each species using single sensillum recordings (SSR), where 86% of the receptors retain the same ligand spectra, while only five OSNs (14%) displayed variation in the odorant that produced the largest response. Responses that were different between the two species are highlighted in orange. (B-E) Here we show immunostaining with neurobiotin (green) and nc82 (red or magenta) across the antennal lobes of *D. suzukii* adults, which were backfilled from the ab2-like or ab3-like sensillum types. Each sensillum type was first identified with electrophysiological contact and odor response recordings from the antenna before application of the neurobiotin. (B) Spatial map of the antennal lobe (AL) atlas of *D. melanogaster*, with corresponding positions (marked in orange) of neurons emanating from the ab2 sensillum. (C) Neuronal wiring in *D. suzukii* that stem from single-sensillum backfills of the ab2-like sensillum, showing similar mapping to the model species. (D) Spatial map of the AL of *D. melanogaster*, with corresponding positions (marked in orange) of neurons emanating from the ab3 sensillum. (E) Neuronal wiring in *D. suzukii* that stem from single-sensillum backfills of the ab3-like sensillum. This spatial congruence between these two species, combined with the morphological (e.g. large basiconic identity) and odor ligand data from the other neuron housed in either ab2 or ab3 sensillum (e.g. Or59b and Or85b), continues to provide strong, corroborating evidence that the wiring for *D. suzukii* and *D. melanogaster* is unchanged for these two sensilla types. This conservation of wiring within the antennal lobe is despite changes in odorant sensitivity and ligand spectra for the ab2B and ab3A OSNs between these two species.

### Conservation of neuronal wiring despite ligand spectrum shifts

We wanted to continue to explore the neural pathways for the ab2-like and ab3-like OSNs in *D. suzukii*, we therefore next documented the circuitry from the antenna towards the antennal lobe (AL) using the single-sensillum backfill neuronal staining technique^29^. By comparing the labeled glomeruli from *D. suzukii* to the known AL atlas of *D. melanogaster* (**Figure 1 B-E**)^30^, we could provide additional support for the ab2-like and ab3-like sensillum type identification, as these two fly species shared identical wiring for these OSNs, despite the observed deviation in ligand spectra for *D. suzukii* adults.

### Spatial mapping of sensillum subtypes

We next mapped the position of all sensillum subtypes on the *D. suzukii* and *D. melanogaster* antenna and palps using several individuals to create an aggregate diagram. The heads of both *D. suzukii* and *D. melanogaster* were positioned in four ways to completely map the spatial pattern for sensillum abundance, including antennal arista down, arista side, arista up as well as across the maxillary palp (**Figure 2A-D**). Here, as we had established ligand spectra for all OSNs of both species, we used SSR data to document the position of each sensillum type on the antenna, which provided spatial information as well as the relative abundance, and this data matches previous studies of *D. melanogaster* antenna^19,31^. The positioning of each specific sensillum type was nearly identical in both species and arranged in concentric circles or zones; however, we did observe large variations in the abundance of sensillum types between species (**Figure 2 E-H**). For example, within the large basiconics, while the ab1 sensillum counts were nearly identical, we found that *D. suzukii* had almost twice as many ab2 as *D. melanogaster* (22 and 12, respectively). In contrast, we found that *D. melanogaster* had more than twice as many ab3 sensilla compared to *D. suzukii* (13 and 6, respectively). It was possible to identify large basiconics due to physical metrics, like width, tip shape and length (**Supplementary Figure 1**). In addition, we could also positively identify large basiconics based on the amplitude of SSR spike response (**Figure 2 I-K**). Here the ab2 and ab3 sensilla are also uniquely identifiable based on the ratio of the A neuron to B neuron spike sizes, where ab2 has an exaggerated disparity between the two neurons, while the ab3 neurons are much closer in spike size (**Figure 2 I-K**). Moreover, the ab1 sensillum is straightforward to identify due to its unique housing of four OSNs, as well as the characteristic response to CO_2_ odor cues^32^. While our sensillum counts are not the absolute total number of large basiconics, this was provided previously for these two species^1^, and we are confident that the preparations and the zones of interest we counted from were the same during the comparison of the two species^25^. As such, these counts represent strong relative values for species-specific comparisons of sensillar abundance. In addition, it is important to note that the flies do differ in absolute size, as does their antennal surface area^1^. However, we still show a consistent difference that is not explained by the larger size of *D. suzukii* relative to *D. melanogaster* adults, therefore, additional evolutionary factors are in play concerning sensillum abundance.

**Figure 2.**
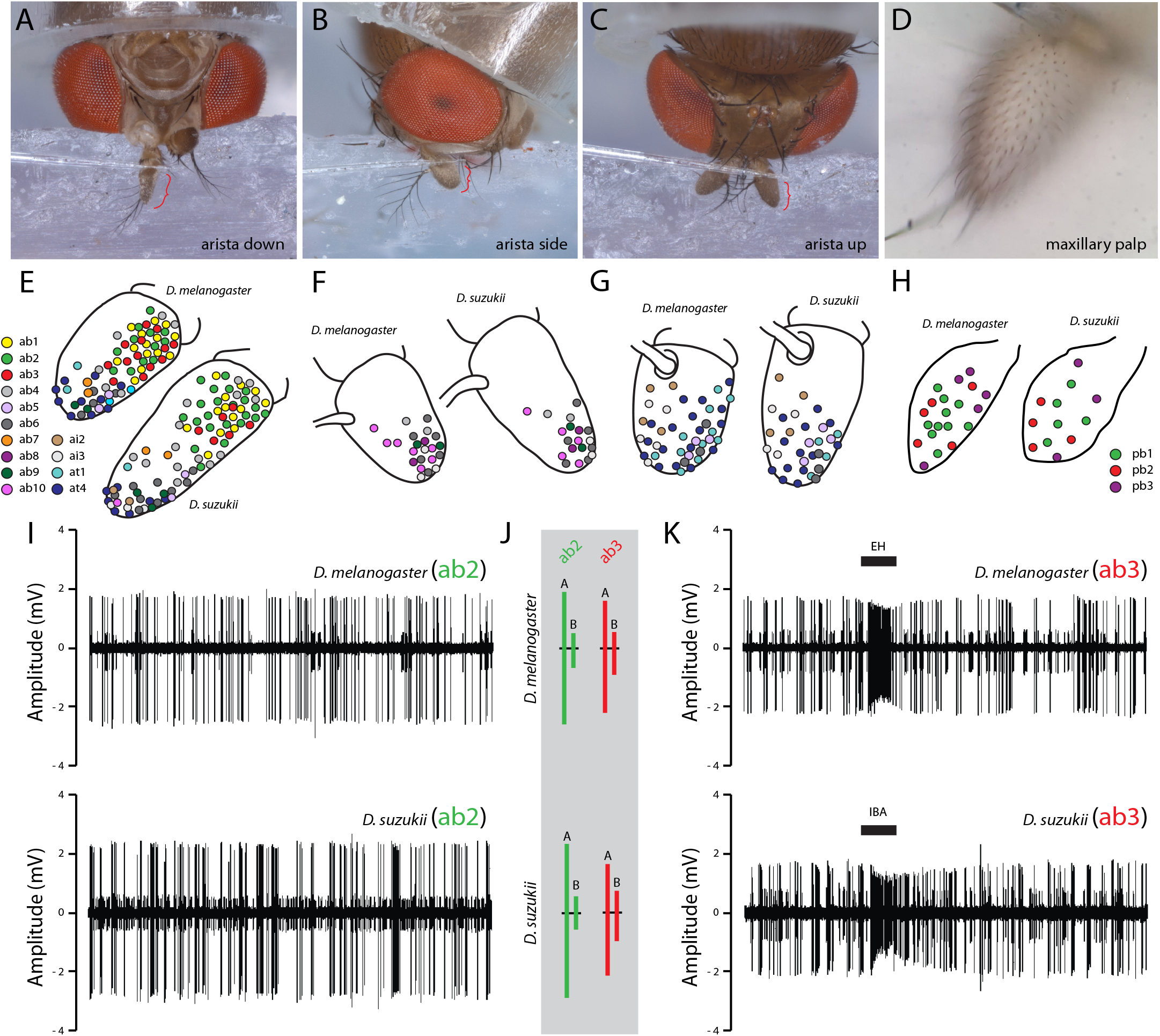
Olfactory sensory neuron mapping across the antennae of *Drosophila suzukii*. (A-D) The antennae of each *Drosophila* species were prepared in several different positions to optimize access to the various sensillar subtypes, including the four positions for *D. suzukii* that are shown. (E) Schematic drawing of the third antennal segment from both species, where spatial distribution and sensillum abundance were identified during single sensillum recordings (SSR). Each color denotes a unique subtype. Shown are the sensillar mappings from the arista down configuration from both target species. (F) Arista side configuration. (G) Arista up preparation. (H) Maxillary palp preparation. (I) Shown are example traces of the ab2 sensillum type, here we note maximum amplitudes well over 2 mV for each species. (J) Consistent ratio of amplitude differences between the A and B neurons for both ab2 and ab3 sensillum. Here we observed that the ratio for A:B in ab2 was larger than for ab3 recordings. We also note that ab3 has smaller amplitudes than ab2 types. (K) Shown are example traces of the ab3 sensillum type, where we note maximum amplitudes at or near 2 mV for each species. Here the relative size of the ab3B neuron amplitude is much larger when compared to ab2B, in relation to their respective A neurons for ab2 and ab3 sensillum types. In general, these stereotyped response dynamics appear conserved between species, and add an additional layer of confirmation to the chemical and morphological identification of sensillum type in these novel *Drosophila* species.

### Proportional analyses of sensilla across *Sophophora*

As we were interested in the evolution of olfaction in *D. suzukii*, we next sought to examine 18 additional species within the *Sophophora* subgenus in order to generate an evolutionary framework of sensillar variation (**Figure 3**). Here we screened each new species for ab1, ab2 and ab3-like sensilla using a wide panel of odorants. The total sample sizes across individuals (n) and across large basiconic sensillum recordings (R) is listed above each species (**Figure 3**), and we present the data via a proportion of these total SSR basiconic contacts. As has been shown previously^19^, we confirm the overrepresentation of the ab3 sensillum for the entirety of the *melanogaster* clade (**Figure 3;** ab3 bias; shown in red), including the confirmation of the most heavily ab3-biased species, *D. sechellia* and *D. erecta*. Intriguingly, we also document that each of the spotted wing species within the *suzukii* clade all display, in contrast to the *melanogaster* clade, a reduction in ab3 and a corresponding overrepresentation of the ab2-like sensillum (**Figure 3;** ab2 bias; shown in green). We believe the ab3 bias of the *melanogaster* clade might be linked to ecologically associated detection of strong fermentation components, while the ab2 sensilla is perhaps more tightly associated with fresh fruit esters or the fruit ripening process, but more work is needed to examine this in behavioral trials. Our screen of the *Sophophora* subgenus also provided evidence for an ab1 bias in several species (**Figure 3;** shown in yellow). We do not have enough ecological information about many of these species, but we can speculate that perhaps there exists a common association with higher altitudes and altered CO_2_ concentrations in these alpine habitats (e.g. *D. birchii)*^33^, or that floral-, foliage-, and yeast-derived host signals^34–36^ might play a role in this association with CO_2_ sensitivity. Lastly, there were four species that had a roughly even proportion of basiconic sensillum types, which are shown in grey. Again, we are greatly limited due to the unknown ecology of most species, thus it is currently unclear what the evolutionary or ecological rationale is for these differences in sensillum abundance.

**Figure 3.**
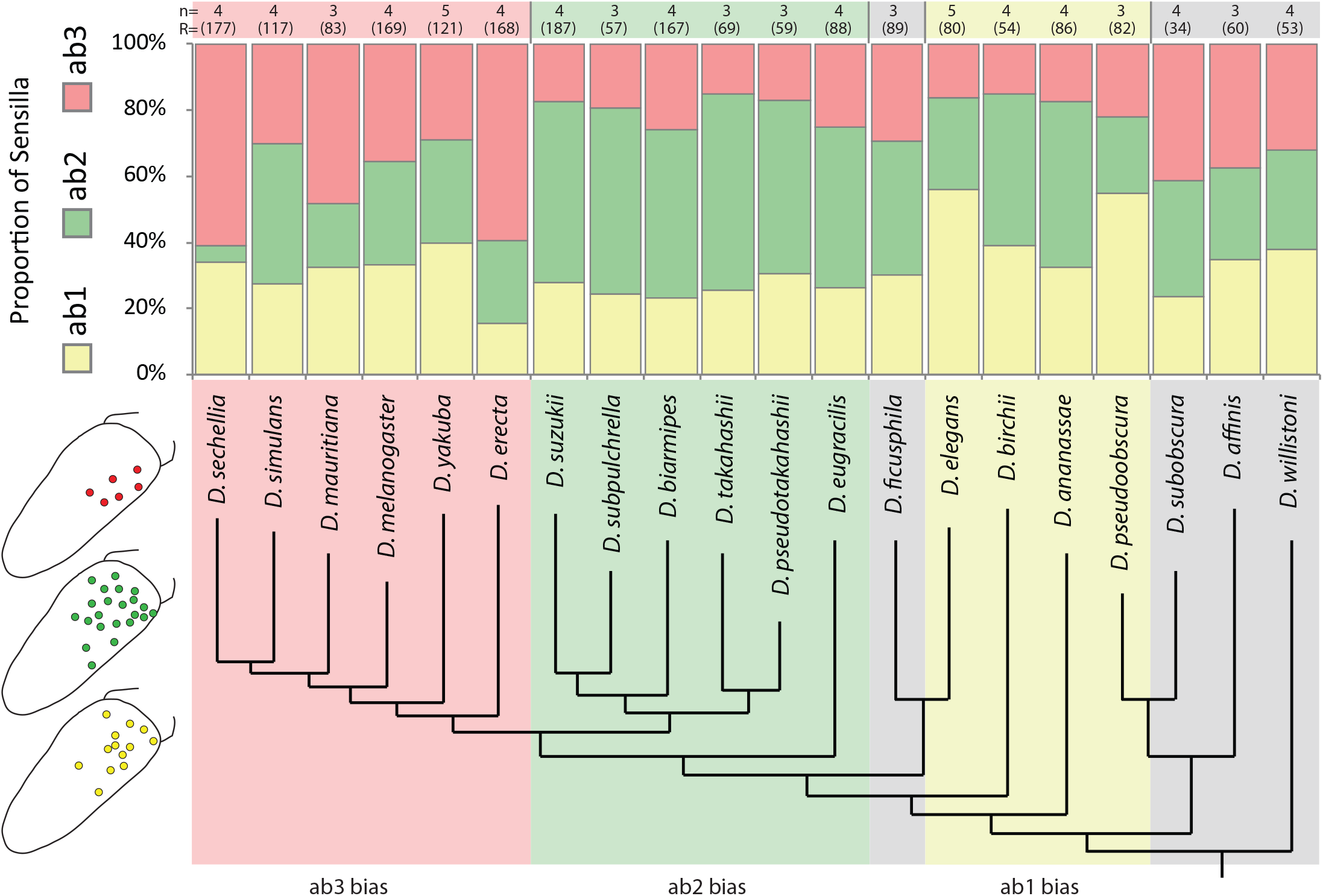
Large basiconic sensilla across the subgenus *Sophophora*. The proportions of ab1 (yellow), ab2 (green) and ab3 (red) sensilla are shown for each of the 20 species examined, with total sample sizes of large basiconic sensillum contacts listed in parentheses above each species, as well as sample sizes. In general, there is a biased, overrepresentation of the ab3 sensillum type for the *melanogaster* clade (those in red on phylogeny), with extreme examples including *D. sechellia*and *D. erecta*, where we also note a drastic reduction in the ab2 sensillum type that is inversely related with the larger proportions of the ab3 type. In contrast, we notice a biased, overrepresentation of the ab2 sensillum type across the spotted wing species, including the *suzukii* clade (those in green on phylogeny). Here, we see a relative reduction in ab3 that correlates with the increases for ab2 sensillum number. In addition (shown in yellow on phylogeny), there is a trend for increased ab1 representation in several species, while those species in grey denote a more even proportion for each of the three large basiconic types.

### Functional ligand spectra for sensillum types across *Sophophora*

As we had established that some of the main olfactory differences between *D. suzukii* and *D. melanogaster* were related to large basiconics, we sought to test this hypothesis by looking at olfactory ligand variation across our 20 species within the *Sophophora* subgenus (**Figure 4**). Here we screened each species with a diverse panel of odorants, including high-throughput testing that utilized fruit, foliage and floral headspace extracts via gas-chromatography mass-spectrometry combined with single-sensillum recordings (GC-SSR). Of the ten OSN types we examined from each of the 20 species, only three showed any dynamic variation in ligand spectra or ligand sensitivities (i.e. ab1C, ab2B, ab3A; **Figure 4**). We note that the ab1C neuron (which co-expresses Gr21a and Gr63a in *D. melanogaster)* displayed a large variation in sensitivity towards CO_2_ between our species, which has been suggested previously^10,37–39^; however, the best ligand for this receptor was found to be identical between all those tested, and ab1C appears to remain narrowly tuned (**Figure 4 B**). Other neurons within this sensillum type, such as ab1A (which expresses Or42b in *D. melanogaster)*, were functionally identical in each new species that we tested, with all species responding most strongly to ethyl acetate. However, two OSNs were consistently variable over the *Sophophora* subgenus, including the same neurons that we originally show were different in *D. suzukii*, namely ab2B and ab3A (**Figure 4 C,D**).

**Figure 4.**
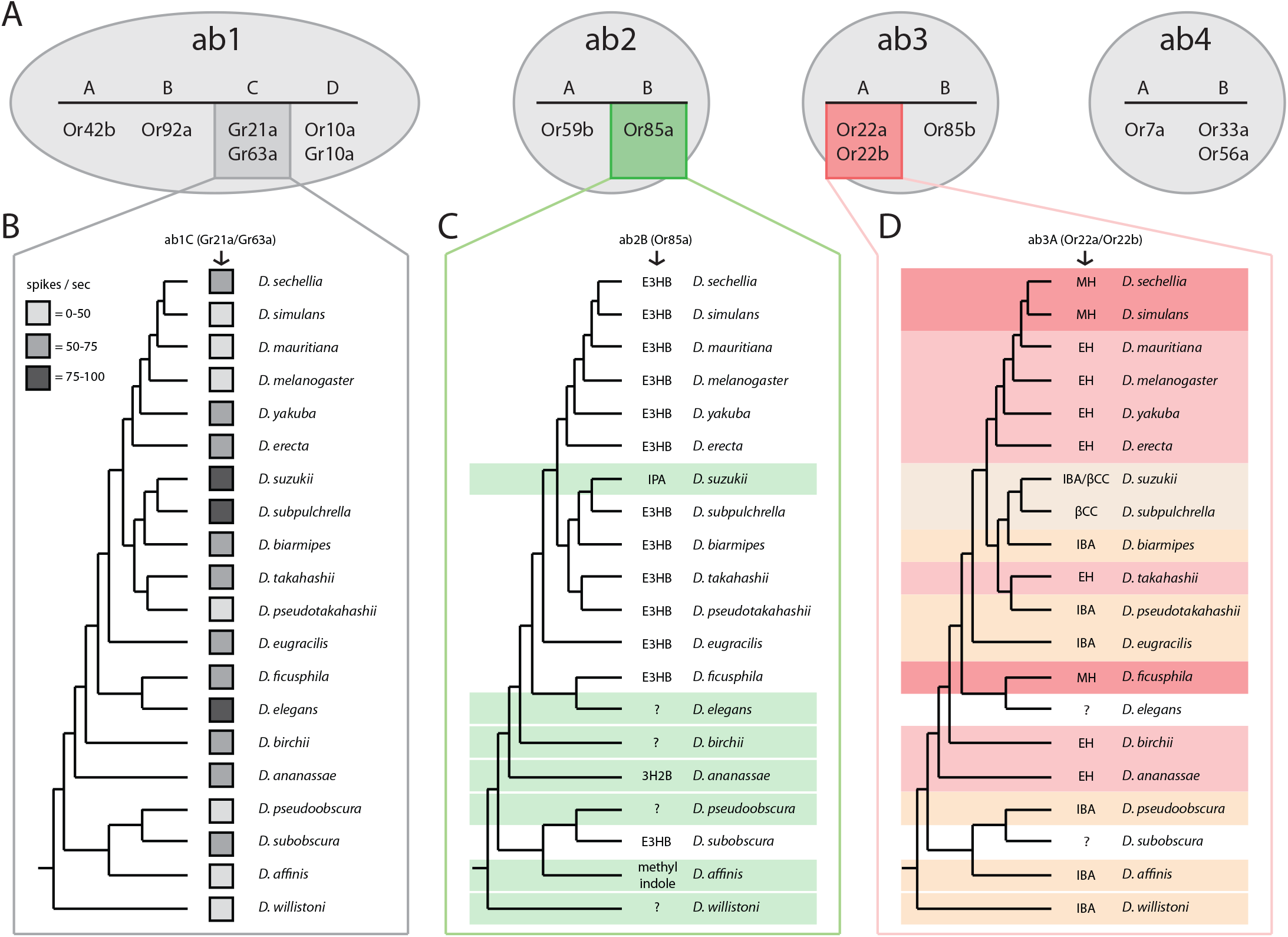
SSR and ligand spectra of 4 sensillum types across 20 species. **(A)** Olfactory receptor neurons (OSNs) shown at top without color were identical in their ligand responses across all 20 tested species (i.e. ab1A which houses Or42b in *D. melanogaster*). **(B)** There was a strong sensitivity variation noted for the ab1C neuron, which contains the CO_2_ detecting GRs; however, the best ligand was identical for all tested species. Additional testing would be required to confirm that CO_2_ delivery was identically administered, and whether this represents a functional sensitivity change between species. **(C)** We note a consistent ligand shift in the ab2B neuron (shown in green, which contains Or85a in *D. melanogaster*). Here the majority of species retained the exact same ligand spectra; however, 7 of the 20 species had an entirely novel ligand, which did not match with any of the other tested species, and we could not identify any odorant that activated this OSN in 4 different species. **(D)** We noted the most consistent changes in ab3A neurons between these 20 species (shown in red, which co-expresses Or22a and Or22b in *D. melanogaster*). Here we observed four or more separate ligand tunings, as defined by the strongest response at the 10^−4^ odorant concentration (diluted in hexane). For two species, we could not identify a strong ligand for this OSN, despite screening with over 80 synthetic compounds and thousands of natural compounds via GC-SSR high-throughput examination using a variety of plant, flower or fruit headspace materials. (E3HB = ethyl-3-hydroxy butyrate; IPA = isopentyl acetate; 3H2B = 3-hydroxy-2-butanone; MH = methyl hexanoate; EH = ethyl hexanoate; IBA = isobutyl acetate; βCC = beta-cyclocitral).

In general, ab3A (which co-expresses both Or22a and Or22b in *D. melanogaster*) was the most commonly changed OSN within our 20 fly species (**Figure 4**), and the response usually fell into one of four main categories of best odorant profile (i.e. methyl hexanoate (MH), ethyl hexanoate (EH), isobutyl acetate (IBA) or beta-cyclocitral (βCC)). The different ligands for ab3A often seemed to be conserved between closely related species. Again, we noted two different functional ab3A types in *D. suzukii*, where one was tuned to IBA (type i) and the other type was tuned best towards βCC (type ii) (**Supplementary Figure 2 A-D**). We observed these two types of ab3-like sensilla within the same individual animal, with a roughly 60:40 split between those detecting IBA and βCC, respectively. Further work is needed to confirm and reconstruct the neural circuit for these two sensillum types in *D. suzukii*, as while they map to the same glomerulus (DM2-like), it was unclear from our neuronal backfills whether type i and type ii map to different regions within this same glomerulus. We note that several functional types of this ab3A OSN have been recently reported within *D. melanogaster* populations^40,41^. One variant expression is a chimeric gene, Or22ab, which arises from the fusion of both the Or22a and the Or22b genes into a single receptor protein^40,41^. Moreover, we note that the ligand spectrum reported for this Or22ab variant is functionally similar to our current data in cases where we observe that IBA is the best ligand for many of these *Sophophora* species, including *D. suzukii* adults. Thus, it is possible that this variant receptor type also occurs within non-*melanogaster* members of this genus. Another consideration is that Or22c, which is usually expressed only in larvae, detects 2-acetylpyridine which bears aromatic, chemical similarities to βCC^26^; therefore, another chimeric form, such as Or22ac, may also be possible in nature (**Supplementary Figure 2 B-E**). Overall, the olfactory changes for ab3A were more of a gradient or moderate shift between species, such as the transition from EH to MH as the best ligand (e.g. *D. melanogaster* and *D. sechellia)*. We also note two species for which we could not identify any strong ligand for ab3A, *D. elegans* and *D. subobscura*. Here, not much is known about the ecology, and despite efforts to screen a variety of headspace collections from flowers, fruits and tree sap (including in total several thousand compounds), we could not identify any suitable ligand candidates for this OSN in these two species.

Intriguingly, we also observed several changes in the ab2B neuron across our 20 examined species. In this case, most species displayed exactly the same ligand (i.e. ethyl-3-hydroxybutyrate (E3HB)). However, seven *Sophophora* members had an acute change in ligand spectra, where none of these seven species shared any overlap for their novel ligand, making each of these adaptations entirely species-specific. For example, as was published previously^10^, we again demonstrated that *D. suzukii* ab2B has changed ligand spectrum towards the detection of isopentyl (isoamyl) acetate (**Figure 1; Figure 4**), which is an odor associated more with ripe as opposed to overripe fruit. For two of the other six species with changes in this ab2B OSN, we could identify a strong ligand (**Figure 4**), including *D. ananassae* (3-hydroxy-2-butanone; a.k.a. acetoin) and *D. affinis* (methyl indole & α-pinene); however, again, very little is known about the ecology of these two Drosophilids. Despite high-throughput GC-SSR screening of the remaining four species, we did not identify any strong ligand candidates. We believe that screening with host-related odors (once ecological information is known) will most likely lead to the identification of the ligands for all of these *Drosophila* species with functional changes in ab2B, but it is also possible that some of these olfactory receptors are non-functional pseudogenes due to natural mutation in the amino acid sequence. Thus, at this stage, we cannot definitively say whether the variation in ab2B is (a) due to a drastic ligand shift in the receptor expression (i.e. protein sequence variation), (b) due to the replacement with a new or duplicated olfactory receptor^42^, or lastly, (c) whether these changes are the result of non-functional pseudogenes. However, at least in *D. suzukii*, we have shown that the ab2B neuron, despite this acute shift in ligand spectrum, remains a fully functional OSN type. Moreover, that this OSN still maps via the same neural circuit to the same location within the AL (**Figure 1 B-E**), and thus might carry a similar behavioral relevance for *D. suzukii* as that of the *D. melanogaster* model.

### Receptor sequence alignments across the *Sophophora* subgenus

As we had established both functional variation and functional conservation throughout OSNs within the *Sophophora* subgenus, we next sought to examine olfactory receptor (OR) protein sequences as a means to correlate and explain our SSR and odorant response data. Our assumption was that when functional SSR data did not vary between species that we also would not expect to observe any significant changes in OR sequence orthologues. Thus, we first pulled as many sequences from our 10 OSNs (e.g. those olfactory receptors housed (at least in *D. melanogaster*) in ab1, ab2, ab3 and ab4 sensillum) from as many of our 20 species as were publically available in databases such as Flybase and GenBank. The alignment of these orthologues provided a wealth of information in conjunction with our SSR functional data screen. Here we observed that the olfactory receptors that were identical between our 20 species during SSR testing were also nearly identical in amino acid sequence (**Figure 5A**; e.g. Or42b). Although it has been suggested that the geosmin-detecting receptor, Or56a, is the most widely conserved receptor across the genus *Drosophila*^18^, here we observe that other ORs are equally or even more functionally conserved, such as Or42b, which is housed in the ab1A OSN and responds to ethyl acetate (an attractive odorant). For the 13 to 15 *Drosophila* species for which protein sequences were available from genomic data, we found very few changes in most receptors, which echoed our lack of functional shifts in ligand sensitivity or selectivity from our SSR datasets (**Figure 5B**; shown is Or42b, and note high pairwise identity between species, in black). However, other receptors, such as those found in ab3A (i.e. Or22a/ Or22b in *D. melanogaster)*, which were highly variable across species in our SSR data, were also demonstrated here to be highly variable in amino acid sequence (**Figure 5C**; shown is Or22a, and note low pairwise identity between species, in white). The same trend was true for ab2B (which houses Or85a in *D. melanogaster)*, where sequence variance mirrored the diversity in SSR response profiles between our species (**Figure 5A; Supplementary Figure 3-6**). Therefore, in general, it appears that olfactory ligand variation positively correlates with amino acid variation in the corresponding OSNs.

**Figure 5.**
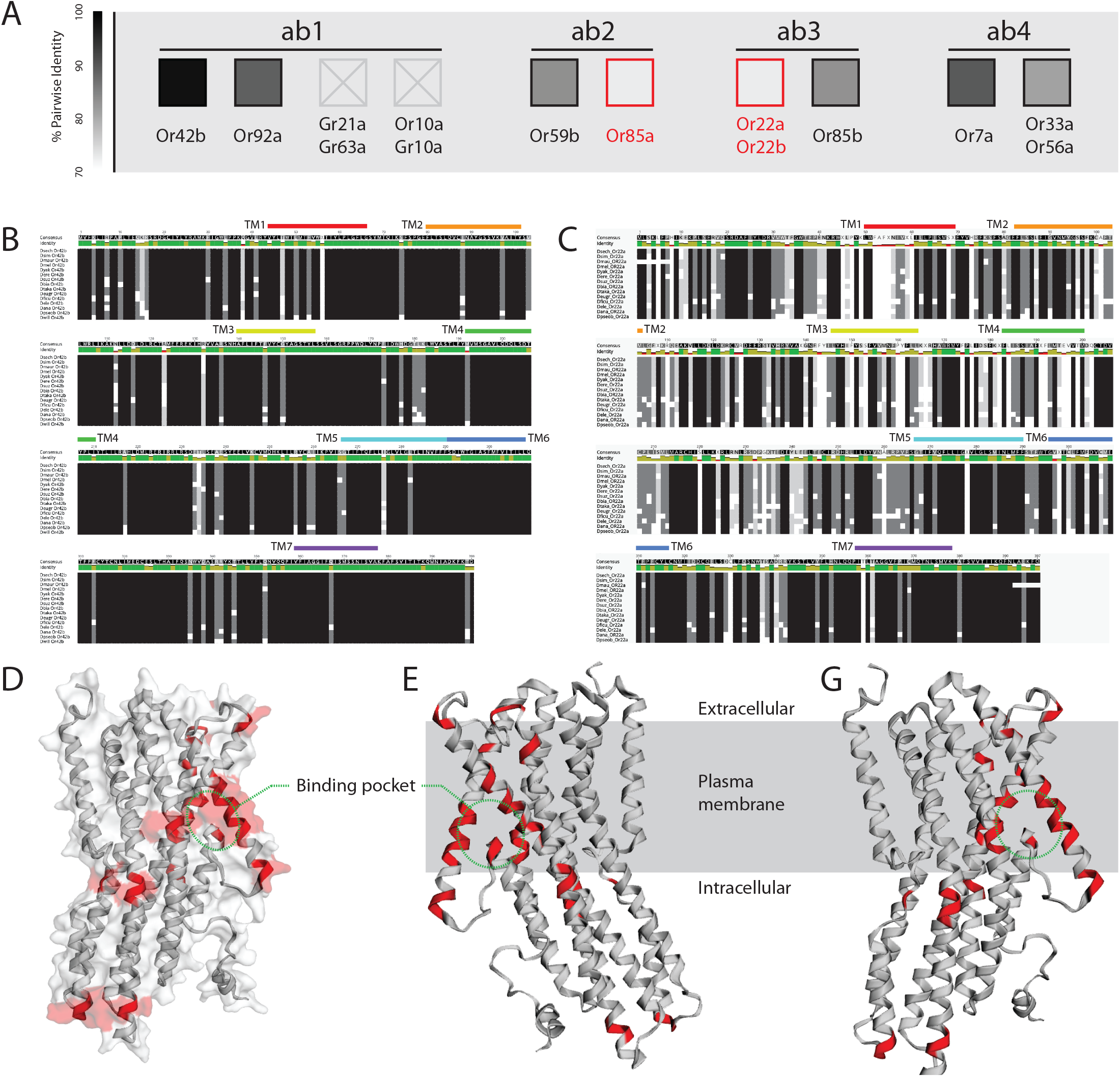
Protein sequence alignments of receptor orthologues. **(A)** Heat map (greyscale) between the sequences representing the percent pairwise identity or matching across the examined species. Darker colors illustrate higher degrees of sequence similarity, and lighter colors denote receptor proteins with high variability in sequence data between *Sophophora* species. Boxes with an X denote receptors with insufficient sequence data available for comparison. In general, most of these eight olfactory receptors were highly conserved across all examined species; however, both Or22a and Or85a showed large variability in protein coding sequences, which mirrors electrophysiological variation in these same receptors (see Figure 4). **(B)** Example of protein alignment for Or42b (ab1A), which was highly conserved across all available species (larger panels available in supplementary files). Note the high amount of black squares that illustrate identical amino acids across fly species. This data strongly matches the identical functional ligand spectra observed for this olfactory sensory neuron, which detected ethyl acetate in all 20 species. Also shown are each of the 7 predicted transmembrane domains (TM1-7). **(C)** Example of protein sequence alignment for Or22a, which has a large variation in amino acid sequence for which data is available between the species (larger panels available in supplementary files). Note the high amount of grey and white squares that illustrate highly variable amino acids across these 15 fly species. This Or22a sequence data strongly supports the quite variable odorant ligand spectra observed for this OSN in recordings from the antenna. Additional protein alignments for the other six receptors are available in the supplementary materials provided with the online version of this publication. Also shown are each of the 7 predicted transmembrane domains (TM1-7). **(D)** Predicted protein folding and 3D surface area for Or22a sequence (397 amino acids), with variations between species highlighted in red. **(E)** Side view (through plasma membrane) of Or22a protein structure, showing changes between species in red. **(F)** Alternate side view with approximate location of plasma membrane relative to the protein tertiary structure. Indicated in red are sequence locations that deviate across all available *Drosophila* species data, including amino acid positions: 45-47, 52-60, 100-101, 110, 143-144, 160-161, 166, 191-192, 196-197, 199, 202, 359.

Moreover, we found that our assumption was correct, that functional SSR variation in ligand spectra matched changes in orthologue sequence data; similarly, that when functional SSR data was identical between species that we did not see sequence changes between those species (**Supplementary Figure 7; Supplementary Figure 8**). The species-specific responses of these two sensillum types (ab3A and ab2B) also show up in a principal component analysis (PCA), with *D. suzukii* closely clustering with *D. eugracilis, D. pseudoobscura*, and *D. biarmipes* regarding the ab3A responses (**Supplementary Figure 10 A**). Likewise, species whose ab3A-like sensilla exhibited similar SSR responses had also a higher similarity regarding their Or22a sequences (**Supplementary Figure 7,8,10**). Here we found the same correlation between pairwise SSR similarities and pairwise sequence similarities for ab2B-like responses and the corresponding Or85a sequences as well. However, *D. suzukii* showed overall much less similarity regarding both SSR and sequence data as compared to the other species (red dots; **Supplementary Figure 10 C,D**), which indicates that Or85a (ab2B) may have been replaced by another receptor protein^42^. Notably, we identified the lowest sequence similarities within the transmembrane regions 1 and 3 (**Supplementary Figure 10 E,F**), suggesting that these regions of the protein might be involved in speciation events across the *Drosophila* genus, as was previously suggested from analyses of the *melanogaster* clade^15^.

However, it continues to be unclear which protein sequence changes are the most likely to be associated with functional olfactory changes. For example, which amino acid substitutions are the most critical and correlate the most strongly with functional odorant profile types. Recently, Auer et al.^15^, suggested that several positions are crucially important for the ligand spectrum shifts observed between the Or22a variants found within the *melanogaster* clade (e.g. between *D. sechellia* which detects MH, and *D. melanogaster* which is tuned more towards EH). This includes a newly described putative binding pocket position for Or22a, which Auer et al.^15^ suggested to be the amino acids found in positions 45, 67, and 93 (from the total length of 397 amino acids for this receptor). With this in mind, we sought to examine our sequence variation and ligand shifts with this new information as a potential guideline, focusing our efforts here on just the Or22a receptor between our 20 species. As such, we examined the predicted protein folding and tertiary conformational changes to the protein when we include the species-specific changes to amino acids (**Figure 5; Supplementary Figure 9**), and looked for a correlation between 3D structure and similar ligand detection in our panel of 20 fly species.

### Protein folding and tertiary structure

Using the well-studied Or22a sequence and the predicted tertiary protein structure from *D. melanogaster*, we compared the locations of the observed variations in sequence data from all available fly species for this receptor (**Figure 5 D-G**). Here we highlighted amino acid positions in red that were variable between our diverse phylogeny of *Drosophila* species (**Figure 5 D-G**), and noted a significant overlap in this positional variation. Moreover, we found that a similar transmembrane region is most commonly changed within the *Sophophora* subgenus (**Figure 5G; Supplementary Figure 9**). This protein structure data supports the changes that have been reported from several members of the *melanogaster* clade^15,20^, as here we found again the same transmembrane regions of interest that were previously identified, and which were predicted to be the putative binding pocket of the Or22a receptor (**Figure 5 D-G,** in red). However, it is not clear how these protein sequence alterations would affect protein folding and thus tertiary structure, therefore additional study is needed to address these hypotheses, especially as it relates to ligand selectivity and binding pocket function^43,44^. As the cryo-EM structure of Orco has now been elucidated^45^, we anticipate a highly researched olfactory receptor such as Or22a would be the next viable candidate for crystallography and cryo-EM studies, and thus would allow for the testing of the before-mentioned hypotheses about functional changes to binding pocket structure. In the present study we also addressed the protein sequences for the gustatory receptors that detect carbon dioxide (CO_2_; Gr21a & Gr63a; ab1C); however, as very few of the target species had available genomic data, it is unclear how these minor changes in sequences account for shifts in ligand sensitivity (**Figure 4B; Supplementary Figure 12**).

## DISCUSSION

Since its introduction into Europe in 2008, *D. suzukii* has been extensively studied, primarily as an agricultural pest, but also as an emerging model for speciation and evolutionary neuroethology^1,10,11,21,46^. Here we provide significant progress towards the functional characterization of all OSNs in both the antenna and the palps of the adult fly. Surprisingly, 86% of the 37 OSNs that we compare between *D. suzukii* and the classical model, *D. melanogaster*, show conserved odorant binding affinities. Thus only five OSNs deviate strongly in their olfactory function, including ab2B, ab3A, ab9B, ab10A, and ai3A, where several of these receptors in *D. suzukii* were predicted to be functionally divergent, according to the analyses of genomic data^47^. For this study, we primarily focus on the two large basiconic sensillum types, and further document that although their odorant tuning differs, *D. suzukii* appears to maintain the same neural connectivity to the primary olfactory processing centers within the antennal lobe (AL) (**Figure 1 B-E**). However, it was shown recently in *D. simulans* that even when peripheral odorant spectra are the same, that changes in neural circuitry can occur in the higher brain centers, and lead to behavioral valence shifts^16^. Thus, additional work should still be conducted to assess all the other OSN connections in *D. suzukii* for variations that may not be apparent at the periphery or within the AL, including behavior.

Another interesting phenomenon that we observe is that there are seemingly two different functional types of the ab3A neuron in this pest insect, one that is tuned towards IBA and another towards βCC, where both types are found in the same animal. This suggests that sensillum types across the antenna are non-uniform, and this may play a wider role in evolution, especially in regard to gene duplication events (e.g. Or22a), where receptor copies may be differentially expressed across the antenna and provide fodder for natural selection to occur and promote disparities in ecological preferences. A similar duplication was shown previously to occur with the Ir75 complex^48^, in this case within *D. melanogaster*, where these co-expressed OSNs map to the same glomerulus, but to differing regions or spatial locations. While our two ab3A OSN types appear to map to the same glomerulus, it is unclear from our study whether they map to different regions within this single glomerulus. Future studies are needed to test the hypothesis that a single glomerulus can split into two or more during the evolution of new odorant function at the periphery^48,49^. In addition, future examination of this phenomenon could address the order or sequence of events that give rise to new olfactory or ecological niches. However, we note that the ab3B neuron remains identical between these two types of ab3 in *D. suzukii* as well as identical to that recorded from *D. melanogaster* adults (i.e. detecting 2-heptanol). Thus both ab3B (Or85b) and ab2A (Or59b) can act as anchors for identifying ab2-like and ab3-like sensilla within all 20 examined *Drosophila*, as these OSNs are highly conserved and seemingly identical throughout our *Sophophora* species despite the paired OSN in the same sensillum often shifting ligands (**Figure 4; Figure 5**).

In the last few years, increasing evidence has been provided to support the notion that the closest relative for *D. suzukii* is actually *D. subpulchrella*, and not *D. biarmipes*^47,50^. Here we identify for the first time a second species which responds strongly to beta-cyclocitral, namely *D. subpulchrella* **(Figure 4**), which further supports the connection. Although the ecology of *D. subpulchrella* is understudied, based on the serrated ovipositor (**Supplementary Figure 11**) it has been assumed that it also lays eggs in fresh or ripening fruit resources^51,52^, just like the *D. suzukii* pest. Besides the similarity of odorant responses across the species members of the *suzukii* clade, we also note for the first time an overrepresentation of the ab2-like sensillum within this genus (**Figure 3**), which is in contrast to the well-studied *melanogaster* clade, which instead have a general overrepresentation of the ab3-like sensillum. As such, we continue to compile additional strong support for the notion that relative size and energy allocation towards a particular odorant is indicative of ecological relevance^1,20,21^, where it appears that *D. suzukii* and other spotted-wing relatives have enhanced their fresh fruit detecting OSNs at the cost of those OSNs that detect fermentation byproducts (**Figure 3**). Moreover, we also show an additional shift in ligand spectra for two fermentation-related neurons, ab2B and ab3A, which in turn now detect ripening fruit odors (**Figure 1; Figure 4**), thus seemingly pushing the *suzukii* clade further towards an ecological niche and host preference that is different from the *melanogaster* clade.

As more energy is devoted towards research into the relatives of *D. melanogaster*, an assumption has been made that this species group is an ecological model for the entire *Drosophila* genus. However, current data illustrates that the *melanogaster* clade is in fact the most divergent from the other members of the *Sophophora* subgenus, where most of the tested species, as well as the more basal species in the phylogeny, show an ab3A odorant tuning towards IBA and not EH or MH odorants (**Figure 4**). Here we also note that most basal species also have different ligands for the ab2B OSN (**Figure 4**). Thus, our data mirror the hypothesis that the evolutionary adaptation of the *melanogaster* clade towards human commensalism and a cosmopolitan lifestyle, which is built around fallen, fermenting fruit, perhaps realted to human agriculture, is purported to be a more recently derived phenotype for this subgroup^41^. Similarly, it is likely that the ecological preference for fermented, as opposed to ripe fruit, has perhaps evolved multiple times across the *Sophophora* subgenus, given that we observe odorant tuning towards EH and MH several times within our screen of 20 species (**Figure 4**). Moreover, we note this potential fermenting fruit preference in *D. ananassae, D. birchii, D. ficusphila*, and again for *D. takahashii*, in addition to the six members of the *melanogaster* clade that we examine. At least for *D. takahashii*, a member of the *suzukii* clade, this fermentation preference was tested and confirmed in previous behavioral trials, where this species preferred to oviposit in fermented strawberries as opposed to ripe fruits^11^. In addition, *D. takahashii* does not appear to have a heavily sclerotized ovipositor, which would be necessary to attack ripe fruit. To our knowledge, this ecological preference for oviposition has not been tested in any of the other species that detect EH or MH via the ab3A OSN type. As such, additional study is required to continue to test our hypotheses about the ecological rationales for ligand spectra shifts in these other understudied *Drosophila* species, but we feel the evidence we provide supports the idea that ab3A (Or22a/Or22b in *D. melanogaster)* is strongly associated with host choice, either for feeding or perhaps indirectly for oviposition preferences as well.

In addition to the changes documented for the ab3A-like OSN, we also demonstrate a series of changes throughout our 20 *Sophophora* species for the ab2B-like OSN. In *D. melanogaster*, this OSN expresses Or85a, and while we find 13 of the 20 species retain the same ligand as *D. melanogaster* (e.g. E3HB), we also uncover seven species with widely varied ligand spectra for this OSN (**Figure 4**). Recently, it was predicted for *D. suzukii* that Or85a was lost during evolution, perhaps due to pseudogenization^47,50^. However, we clearly demonstrate within our current study that a fully functional OSN is present in ab2B (**Figure 1; Figure 4**). Thus, several options exist to explain this occurrence. First, it is possible that Or85a is a non-functional pseudogene that is not expressed in *D. suzukii* or that this receptor is not transported to the cell membrane by an Orco chaperone. In this case, it would be likely that a different receptor is instead expressed in this OSN location as a replacement, and that this new receptor provides the novel SSR response profile that we describe. In this scenario, the closest olfactory receptor match for the observed ligand spectra in *D. suzukii* would be Or47a from *D. melanogaster*, which is co-expressed with Or33b within the ab5B sensillum subtype. Second, it is also possible that Or85a is only predicted to be a nonfunctional pseudogene from the genomic analyses of *D. suzukii* due to a premature stop codon. In this case, a shortened sequence for Or85a in *D. suzukii* might retain functional expression, albeit with a novel ligand spectrum, and thus Or85a acts as a pseudo-pseudogene, which is a phenomenon that has been previously described in *D. melanogaster*^53^. However, this has only been documented for an ionotropic glutamate receptor (IR), and the present suggestion would be the first known case of an OR that acts as a pseudo-pseudogene. In either case, we demonstrate that a functional receptor exists at this OSN location in *D. suzukii*, and that the ligand spectrum deviates strongly from that found in *D. melanogaster*. Interestingly, we also document this same type of occurrence for ab2B in six other *Drosophila* species from our screen (**Figure 4**), where in each case, the ligand spectrum is entirely species-specific, and does not overlap with any other known species. This is unlike the slow changes or gradual shifts observed in ab3A (**Figure 4**), where olfactory deviations in ligand spectra are often shared across several species. With all this in mind, whatever the cause for this change in the ab2B OSN, we show it occurs repeatedly, and it is therefore likely to provide a novel avenue for rapid olfactory evolution throughout the *Sophophora* subgenus.

Through the previous examination of the olfactory system of several species within the genus *Drosophila*, several mechanisms have been proposed for the evolution of chemosensory receptors. It has been demonstrated that alterations in the neural circuitry of the P1 neurons (or neurons in higher brain centers) can dictate attraction or aversion for different species, despite the conserved peripheral detection of an odorant^16^. There have also been numerous publications describing the net gains or losses of chemosensory receptors that result in dramatic changes to host or habitat preferences, such as the variations shown for the *Scaptomyza* leaf-mining genus, which are still within the Drosophilidae family, but have for example, lost Or22a entirely^54^. In addition, there has also been a plethora of examples describing alterations in the relative abundance of chemosensory gene expression, either increases or decreases, where each change corresponds with ecological specialization in regard to either host or courtship preferences^1,15,20,21^. In these cases, the peripheral abundance also always accompanies a corresponding shift in glomerulus volume within the AL^19,30^. In the present study we provide several novel examples of the latter two of these mechanisms, both by illustrating a ligand spectrum shift at the periphery, and by providing robust evidence for sweeping changes in the relative abundance of OSN types. Intriguingly, we find that these two mechanisms often co-occur, that is, when ligands change their binding affinity, we also notice changes in relative sensillum abundance connected to those same OSN types. As such, it is difficult to determine what changes first in the timeline of olfactory evolution, ligand specificity or receptor expression. Therefore, future studies of *Drosophila* should continue to compare closest relatives or entire clades of species in order to determine the potential chronology of evolution, which may or may not have a consistent temporal mechanism within this genus.

An important observation from our dataset is that some receptors are more likely to change than others, as we find for example, consistent alterations to ab2B and ab3A across our 20 species, while other OSNs remain functionally identical as well as continue to be highly conserved in their amino acid sequence, such as those within ab1 and ab4 sensillum types. One explanation for these specific OSN changes could be related to yeast or host-specific microbial odorants. It has been shown previously for *Drosophila* species that yeast control attraction and oviposition more so than the host plant material^55^, thus it is possible that the consistent evolution of these OSNs, especially ab2B and ab3A, is related to yeast-specific and ecologically vital odors for each fly species. Another possible explanation for the observation that evolution more consistently occurs for a particular OSN could be related to the protein stability of tertiary structures or the stability during folding of the protein sequence itself^56,57^. It is possible that Or22a for example is one of the more unstable tertiary structures of all the olfactory receptors within *D. melanogaster*, thus that inherent structural elements make this olfactory chemoreceptor much more susceptible to evolutionary pressures simply by being more malleable. In a similar fashion, it is also possible that a stability explanation could account for why some amino acid regions within a protein sequence are also more likely to change. For example, why amino acid position 45, 67 or 93 of the sequence for Or22a are more likely to be altered between 6 species within the *melanogaster* clade^15^, something which we also show is the case across our 20 examined species (**Figure 5**). Therefore, future scientific objectives should target questions related to the conflicting evolutionary pressures for maintaining protein stability versus the advantage of an opportunistic ability of a protein to evolve rapidly through a high-likelihood of mutations within an unstable or malleable region of the sequence, such as the potential binding pocket of Or22a across the *Sophophora* subgenus^15^. In addition, future research should continue to analyze protein sequences and tertiary structures involved in chemosensation with comparisons across large panels of functional odorant screenings, as this may generate a stronger predictive model for identifying ligands in novel *Drosophila* species using existing genomic data as guidelines. Moreover, perhaps through the generation of machine-learning algorithms, this strategy may eventually lead to predictive properties for binding pocket structures in relation to their function throughout a wider collection of receptor-ligand combinations, and in turn delineate the mechanisms for olfactory evolution across invertebrate and vertebrate systems.

## CONCLUSIONS

In this study we highlight that understanding the evolution of olfaction requires not just the determination of which chemical odorant is detected by a given OSN, but also how the relative abundance of those detectors combines to generate a more holistic view of olfactory function within the *Drosophila* phylogeny. Here we first generate a complete olfactory map of odors detected by the agricultural pest and neuroethology model, *D. suzukii*, which we place in direct comparison to the molecular genetic model of *D. melanogaster*. During this examination of the olfactory machinery for this pest insect, we observe that the majority of olfactory ligands are conserved across the antenna and palps for the 37 identified OSN types (**Figure 1 A**). Moreover, we identify only five main differences in the OSN response profiles of these species, and subsequently we focus in more depth on two of those OSNs, namely the ab2B and ab3A neurons. Unlike these OSNs in *D. melanogaster*, which detect odors commonly associated with heavily fermented fruit (i.e. ethyl-3-hydroxybutyrate and ethyl hexanoate), we determine that these same OSNs in *D. suzukii* are instead tuned towards more volatile esters, namely those odors that are primarily produced in earlier stages of fruit ripening (i.e. isopentyl and isobutyl acetate). This difference between species has been suggested previously^10^, albeit via a much smaller chemical library and odorant screen than the present study. In addition, we also now document not just this difference in chemical detection, but also the difference in relative abundance of these sensillum types between these species (**Figure 3**).

Next, we expanded our study to encompass a total of 20 species that serve as models for the entire subgenus *Sophophora*. Here, *D. melanogaster* and its sibling species display an overrepresentation of ab3, seemingly at the cost of ab2 abundance, whereas in contrast, *D. suzukii* and its spotted-wing sibling species display the largest representation of ab2, and a decreased allocation of resources towards the ab3 sensillum type (**Figure 2 C-E; Figure 3**). Furthermore, besides the variation or tradeoff between sensillar proportions for these 20 species, we also document best ligands across 10 different OSNs (i.e. those housed within ab1, ab2, ab3 and ab4 sensilla), where we again note that only two of these OSNs show significant odorant variation at the periphery throughout the phylogeny (**Figure 4**). This screen across 20 species is in agreement with what we described between *D. suzukii* and *D. melanogaster*, where the major changes we observe occur only within the ab2B and ab3A OSN types. Lastly, in order to address the variation in chemistry detected by these OSNs, we examine the molecular sequences of eight olfactory receptors for which data is available for many of our *Sophophora* species. Here we describe that receptors with few functional changes in odorant detection also have few changes in their amino acid sequence, and thus a conserved protein structure (**Figure 5; Supplementary Figure 3-6**). However, we also show that for the two most variable OSNs in regard to odorant affinity, namely ab2B (Or85a) and ab3A (Or22a/Or22b), that there is also a large deviation in both amino acid sequence and protein structure (**Figure 5**). Again, the transmembrane region of the protein that we identify as the most variable was also recently reported and predicted to be the putative binding pocket for Or22a^15^. Therefore, it is likely that this region of the protein sequence and corresponding tertiary structure is the most pertinent to continue to examine in regard to how specific amino acid substitutions relate to ligand spectra shifts between species, or perhaps towards protein stability^58^.

However, the persistent dearth of viable ecological and natural history information for the majority of the known *Drosophila* species makes the extrapolation towards evolutionary pressures or niche partitioning difficult^59^. This paucity of ecological and host information leaves several species without known ligands for either ab2B or ab3A, despite a chemically diverse and robust screening of these OSNs in the present study using known host materials from other members of this subgenus. As such, we continue to implore the expansion of scientific research to include a wider array of *non-melanogaster* species, especially those investigations that could provide ecological, host, and environmental, habitat or behavioral rationales for morphological and neuroanotomical variation. We expect that as increased efforts are placed on additional *Drosophila* species, that we may in the future identify and test additional hypotheses or explanations for the observed olfactory shifts within this incredibly diverse genus of flies. Through the utilization of the comparative method to study sensory biology (i.e. auditory, visual, olfactory, gustatory and tactile cues) and to study behavioral ecology (i.e. feeding, ovisposition, attraction and aversion), we can increase our understanding of the fundamental mechanisms by which evolution shapes the nervous system. Moreover, we can begin to understand and explain discrepancies between species across the vast array of behavioral responses and behavioral preferences displayed towards the sensory stimuli that these species encounter in their natural environments.

## Acknowledgements

This research was supported through funding by the Max Planck Society (Max Planck Gesellschaft). Wild-type flies were obtained from the San Diego Drosophila Species Stock Center (now The National Drosophila Species Stock Center, Cornell University), as well as obtained from Ehime University at Matsuyama (EHIME-Fly, Japan), which is the branch laboratory for *Drosophila* resources under the National BioResource Project (now moved to Kyorin University, Japan; KYORIN-Fly). We express our gratitude to S. Trautheim, D. Veit and their teams for their technical support, expertise and guidance at MPI-CE. We also thank V. Grabe, F. Miazzi and S. Das Chakraborty for their suggestions and comments on the project. We especially appreciate the advice and guidance on immuno staining protocols from both R. Steiber and L.L. Prieto-Godino. Additional thanks to J. Rybak and L. Gruber for their oversight and assistance in regards to SEM images.

## Author Contributions

This study was built on an idea conceived by I.W.K., while J.Z., A.D.C., M.K. and B.S.H. all contributed to the design of the study. I.W.K. and A.D.C. handled the immunohistochemistry and antennal lobe backfills plus neuroanatomy. I.W.K. conducted all electrophysiology and antennal mapping of olfactory sensory neurons for each *Drosophila* species. I.W.K., G.F.O. and J.Z., retrieved, annotated and arranged all receptor protein sequences for comparison and analysis. J.Z., G.F.O. and I.W.K. described protein folding and tertiary structural dynamics. M.K. and I.W.K. addressed all statistical assessments, while all figures and drawings were prepared by I.W.K. The original manuscript was written by I.W.K., was subsequently edited by M.K. and B.S.H., and all coauthors contributed to the final version of the paper.

## Supplementary information

All data supporting the findings of this study, including methodology examples, raw images and z-stack scans, molecular sequences, accession numbers, statistical assessments, Excel tables as well as other supplementary materials are all available on request.

## METHODS

### Contact for Reagent and Resource Sharing

Additional information or requests for resources and reagents should be directed towards the senior authors:

> Prof. Dr. Bill S. Hansson
>
> Dr. Markus Knaden

### Insect Rearing and Fly Stocks

Flies were obtained from The National Drosophila Species Stock Center, NDSSC, at Cornell University (Ithaca, USA), or from Ehime University (EHIME-Fly; Matsuyama, Japan). Stock numbers and reference specimens include the following genetic lines: *D. sechellia* (14021-0248.07), *D. simulans* (14021-0251.01), *D. mauritiana* (E-18901), *D. melanogaster* Canton-S (Hansson Lab Strain), *D. yakuba* (14021-0261.38), *D. erecta* (14021-0224.01), *D. suzukii* (14023-0311.01), *D. subpulchrella* (E-15201), *D. biarmipes* (14023-0361.10), *D. takahashii* (E-12201), *D. pseudotakahashii* (E-24401), *D. eugracilis* (E-18101), *D. ficusphila* (E-13301), *D. elegans* (E-13201), *D. birchii* (E-24201), *D. ananassae* (14024-0371.12), *D. pseudoobscura* (14011-0121.00), *D. subobscura* (14011-0131.04), *D. affinis* (14012-0141.00), *D. willistoni* (14030-0811.24). All fly stocks were maintained on standard diet (normal food;^1^) at 22°C with a 12 hr light/dark cycle at 40% humidity. In addition, the following species had their diet supplemented with freshly crushed blueberries: *D. sechellia, D. suzukii, D. subpulchrella, D. biarmipes, D. takahashii, D. pseudotakahashii, D. ficusphila, D. elegans, D. pseudoobscura, D. subobscura* and *D. affinis*. Fly vials were maintained with consistent numbers of founding females (15-20) per container, in order to maintain consistent adult sizes via controlled population density. Phylogenetic information for all species was made available from previous publications^1^, as well as other literature^47,51^.

### Chemical stimuli and single sensillum recordings

All synthetic odorants that were tested were acquired from commercial sources (Sigma, www.sigmaaldrich.com and Bedoukian, www.bedoukian.com) and were of the highest purity available. Stimuli preparation and delivery for electrophysiological experiments followed previously established procedures, and any headspace collection of plant, fruit or microbial volatile odors was carried out according to standard procedures^10,60^. For SSR experiments, 10μl of a dilution (10^−4^ in hexane) of an odor was loaded onto a filter paper disc that was placed inside a glass pipette. Electrophysiological contacts were made with tungsten electrodes (reference electrode into the eye, recording electrode into a single sensillum). Females were used from all species, and flies were between 2 and 7 days post-eclosion. An odor panel of 80 compounds was selected based on previous literature^10,11,18,22,61^, and was used to screen all OSNs across the antenna and palps of each examined species. In addition, when this odor screen failed to identify any strong ligands, we also utilized gas-chromatography single-sensillum recordings (GC-SSR). Here we employed odor collections from diverse floral, fruit, and microbial origins. In total, we estimate our GC-assisted screens included between 3000 and 5000 separate odorants, similar to previous studies^28^.

### Neuronal staining (single-sensillum backfills)

Flies were prepared as usual for SSR inside a plastic pipette tip, and sensilla were identified first with characteristic odor screening using tungsten electrodes. The tungsten recording electrode was then removed and replaced with a pulled glass capillary which contained a filament, where the filament was pre-filled with neurobiotin via dipping the unsharpened end of the capillary into a 1-2% solution of the dye. Contact with the targeted sensillum was re-made on the SSR table with this filled glass electrode, which punctured, but did not pierce through both sides of the sensillum. A good contact was established when viable SSR spikes were observed, and when appropriate odor responses could be generated using this glass electrode (which replaced the tungsten wire). Light illumination was then removed, and this sensillum contact was maintained for 30-45 minutes, with periodic odor puffs to help push the dye towards the antennal lobe (AL). Both the A and B neurons were stimulated with their relevant odors every 5-10 minutes for the duration of the contact with glass-filled electrode and neurobiotin. Thus axonal projections of OSNs from ab2 and ab3 sensilla were identified by SSR followed by this neurobiotin backfill. Sensillum types in *D. suzukii* were identified by SSR with the diagnostic odors IPA/methyl acetate for ab2 and 2-heptanol/IBA for ab3. Next, the recording electrode was replaced with a pulled glass capillary (with filament) that was filled with neurobiotin (Invitrogen, 2% m/v in 0.25 M KCl). Neurobiotin was allowed to diffuse for 45-90 minutes under periodic stimulation with the associated odors for each sensillum type. Brains were then dissected in PBS and fixed in 4% paraformaldehyde (PFA) for 30 min at room temperature (RT), rinsed 3 x 15 min in PBS with 0.3% Triton X-100 (PT). This was followed by incubation with mouse monoclonal NC82 antibody (1:30, CiteAb, A1Z7V1) and streptavidin conjugated with Alexa Fluor 555 (1:500, S32355, Invitrogen) in 4% normal goat serum (NGS,) in PT (48 h at 4 C). Samples were washed 4 x 20 minutes in PT, incubated overnight with Alexa633-conjugated antimouse (1:250, A21052, Invitrogen) in NGS-PT, then rinsed 4 x 20 minutes in PT and mounted in VectaShield (Vector Laboratories)^48^. Images were acquired with a Zeiss 710 NLO confocal microscope using a 40x water immersion objective. The *D. suzukii* DM2, VM5d, DM4 and DM5 glomeruli identity was assigned based on similar 3D glomerular position and shape within the AL, as compared to those of the *D. melanogaster* antennal lobe atlas^30^.

### Mapping and counts of olfactory sensillum types

Adult flies were prepared as has been described previously for single-sensillum recordings^10^. A single adult was immobilized in a plastic pipette tip, with only the head protruding. The fly was positioned in one of four ways, in reference to the arista (e.g. arista down, up, side 1 and side 2) (Figure 2A-D). This positioning allowed for consistent orientation of the sensillar zones along the antenna of each species, and enabled consistent counting of sensillum types across individuals. We observe concentric rings or circular organization of the different sensillum types, especially the large basiconics. In this case, the ab3 is usually in the slight depression of the 3^rd^ antennal segment, followed by round concentric zones that increase in first ab1 and then ab2 sensillum types, and subsequently the state of small basiconics such as ab4. Schematics of each species were produced based on contacts with each sensillum type, where a sensillum was identified using physical metrics (i.e. size, tip shape, width) (Supplementary Figure 1) as well as ligand identities of each OSN, and in addition, the electrical response dynamics such as amplitude and relative ratio of OSN firing rates or size (Figure 2 I-K). We found all 20 species had similar morphological characters for large basiconics, as well as electrical response dynamics; however, the density or abundance of each sensillum type varied greatly between species, but not between individuals. Previous estimates of abundance^19^ utilized a sample size of 30-40 contacts, whereas here, we attempted to make between 40 and 100 contacts on average per species (Figure 3). Our estimates of sensillum proportions match very with previous examinations, which were universally restricted to the *melanogaster* clade, thus all species beyond this group are to our knowledge, newly described here concerning sensillar proportions.

### Fly images (SSR heads, wings and ovipositor)

Dispatched flies were mounted and views of the ovipositor (180x) were acquired as focal stacks on an AXIO Zoom V.16 (ZEISS, Germany, Oberkochen) with a 0.5x PlanApo Z objective (ZEISS, Germany, Oberkochen). The resulting stacks were compiled to extended focus images in Helicon Focus 6 (Helicon Soft, Dominica) using the pyramid method. Here we provided images of the serrated ovipositors of close relatives to *D. suzukii* in order to highlight the physical deviations in egg-laying potential (Supplementary Figure 1), which has been described previously^51^. We also documented the differences in male wing pigmentation (32x), as *D. suzukii* and *D. subpulchrella* can be difficult to distinguish (Supplementary Figure 11). Moreover, *D. subpulchrella* were shown recently in the most up to date phylogenetic analyses of this *Drosophila* clade to be the closest relatives to *D. suzukii*, as opposed to *D. biarmipes*^1,21,47,50^. Images were also compiled of dissected heads (128x) to illustrate the mounting preparations for sensillum counts (Figure 2A-D).

### Statistical assessments and figure generation

All images and drawings are originals, and were prepared by the authors for this publication. Figures were prepared via a combination of Syntech AutoSpike32 (v3.7), R Studio, Microsoft Excel, Adobe Illustrator CS5, EzMol (v1.22), and Geneious Prime (v2019.0.4). Statistics were performed using GraphPad InStat version v3.10 and Past v3.25 at α = 0.05 (*), α = 0.01 (**), and α = 0.001 (***) levels. For transmembrane region predictions, several resources were utilized (TMHMM Server v2.0; http://www.cbs.dtu.dk/services/TMHMM/; TMpred Server – EMBnet; https://embnet.vital-it.ch/software/TMPRED_form.html), and any nucleotides converted to amino acid sequences were performed using ExPASy (SIB Bioinformatics Resource Portal; https://www.expasy.org/).

---

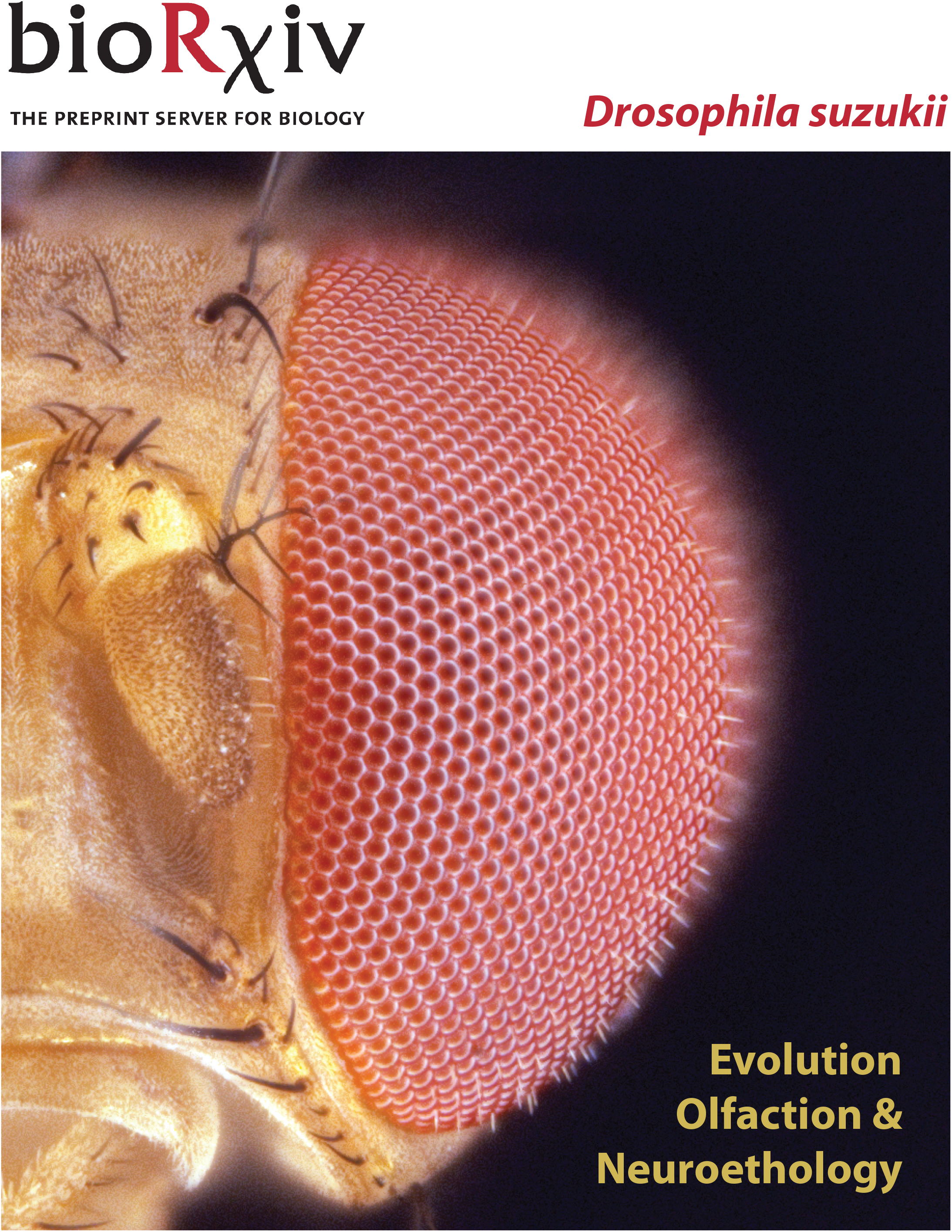

