## Supplementary File 1 for "Evolution of a pest: towards the complete neuroethology of *Drosophila suzukii* and the subgenus *Sophophora*"

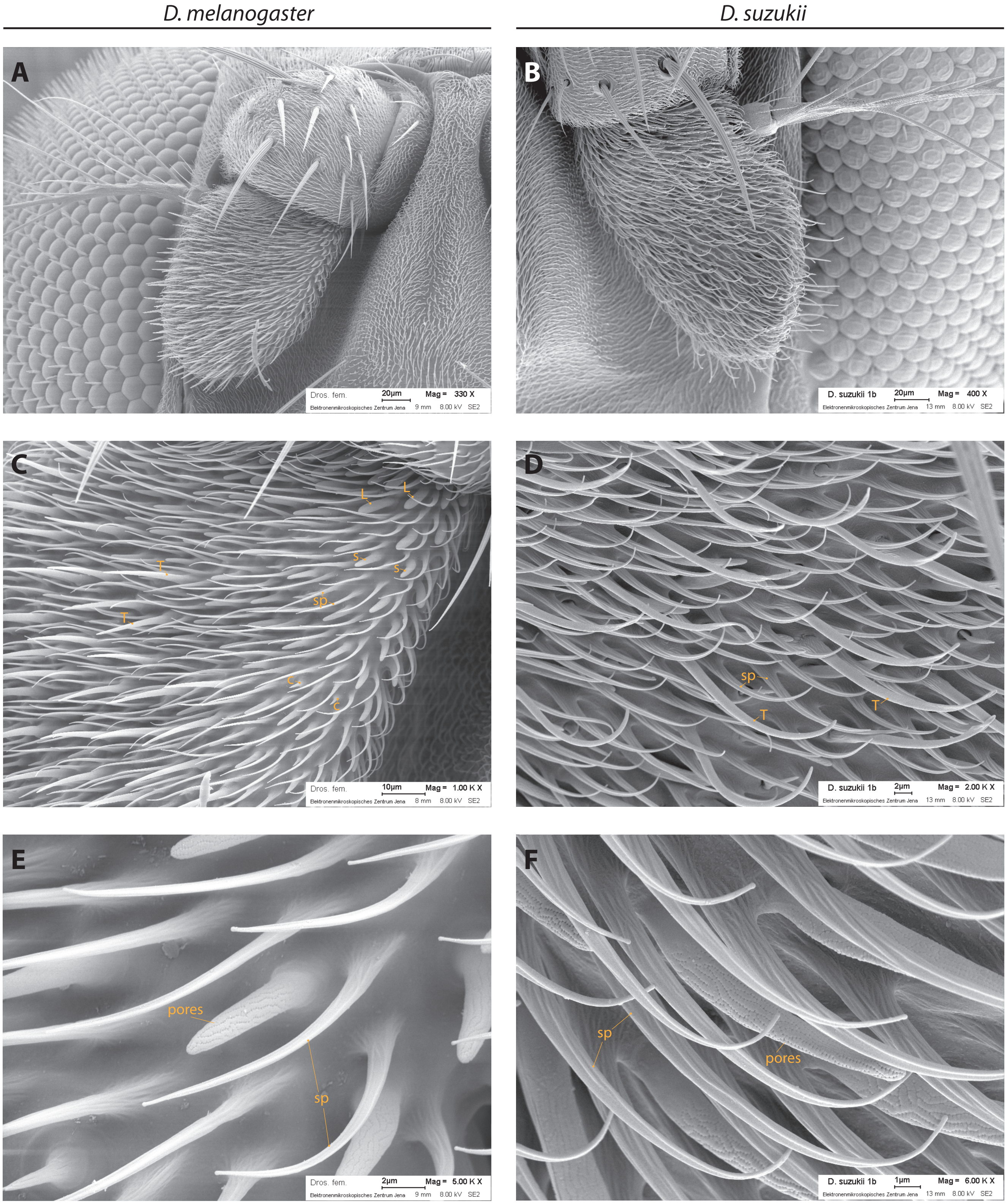


**Supplementary Figure 1. Scanning electron microscopy (SEM) of antenna for *D. melanogaster* and *D. suzukii* adults.**
Comparisons of D. melanogaster and *D. suzukii* antenna, with emphasis on the third antennal segment, the funiculus. **(A)** 330x magnification of the complete antenna of *D. melanogaster* female. Note the numerous straight, short trichoid sensilla that protrude from the antennal surface. **(B)** 400x magnification of the complete antenna of *D. suzukii* female. Note the reduced number of trichoid sensilla, as well as surface area containing these pheromone-detecting subtypes. The trichoids of this species also differ in their increased length, as well as curvature. **(C)** 1000x magnification of *D. melanogaster* antenna, focusing on large and small basiconic sensillum types, which are linear and straight. **(D)** 2000x magnification of *D. suzukii* antenna, where multiple basiconic and trichoid/intermediate sensilla are visible. Note the characteristic curvature. **(E)** 5000x magnification of *D. melanogaster* small basiconic as well as coeloconic sensillum types. **(F)** 6000x magnification of *D. suzukii* basiconics, where curvature is clearly visible, as well as pore structure. (L = large basiconic; s = small basiconic; c = coeloconic; T = trichoid; sp = non-innervated spinule)

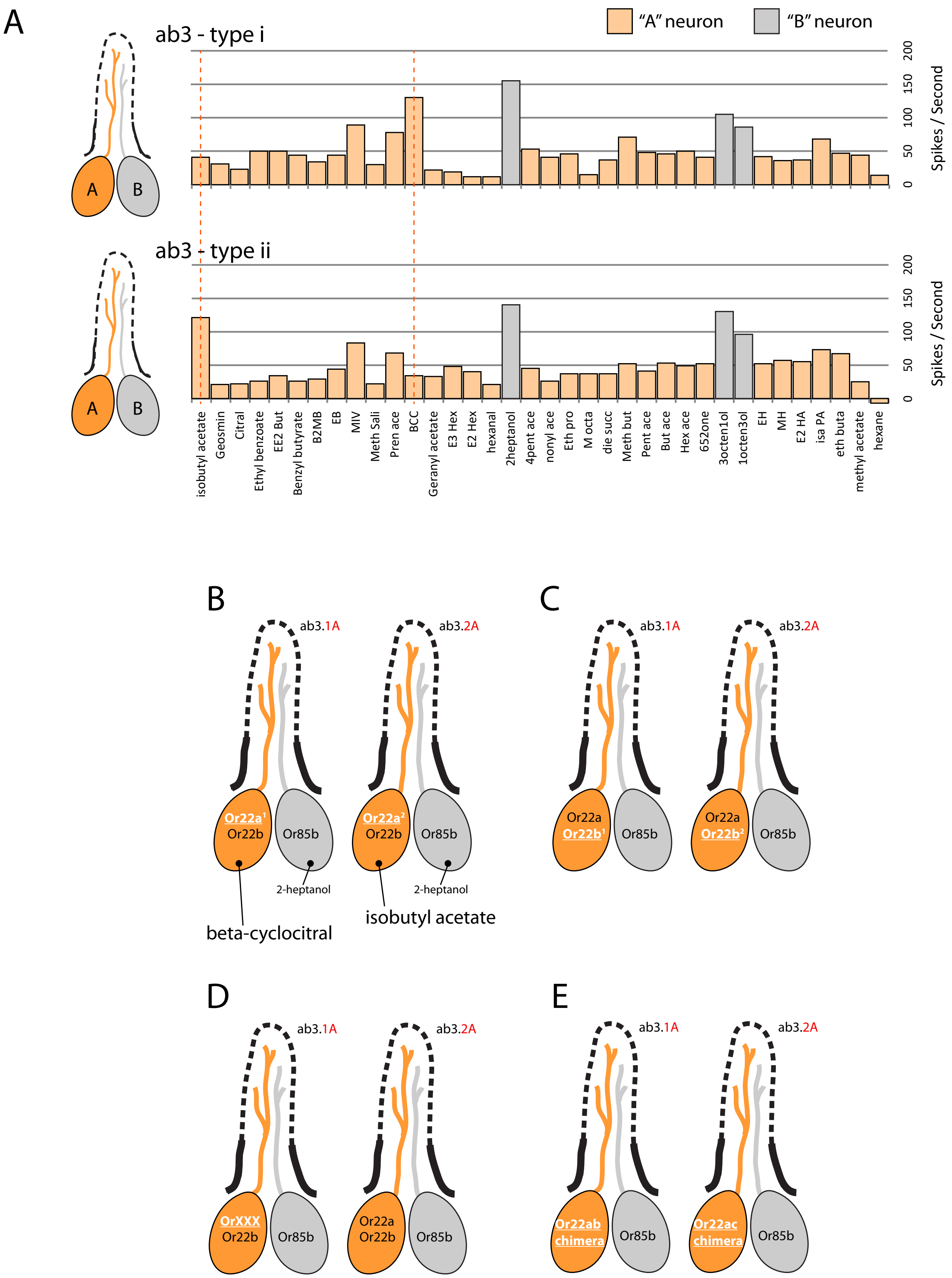


**Supplementary Figure 2. Electrophysiology data for ab1, ab2, ab3 and ab4 sensilla.**
**(A)** Shown are the SSR data (spikes per second) for several odors within the olfactory screen for *Drosophila suzukii* adults. Here we note that the ab3B neuron between type i and type ii ab3 sensilla within the same individual fly are identical, and identical to *D. melanogaster* recordings. However, we also note a ligand shift towards either IBA or βCC, and thus explain the data by proposing two variations of the ab3A OSN across the antenna of this pest species. **(B-E)** Diagram of possible rationales for the evolution of these two types of ab3A observed in D. suzukii adults, where in D. melanogaster this OSN co-expresses two receptors, Or22a and Or22b. Here we include two different functional types of Or22a **(B)**, two different types of Or22b **(C)**, or alternatively, **(D)** the replacement with a novel olfactory receptor. **(E)** Two possible chimeric forms or Or22a, one resulting from a fusion with Or22b, and a second option as a fusion with Or22c (larval receptor).

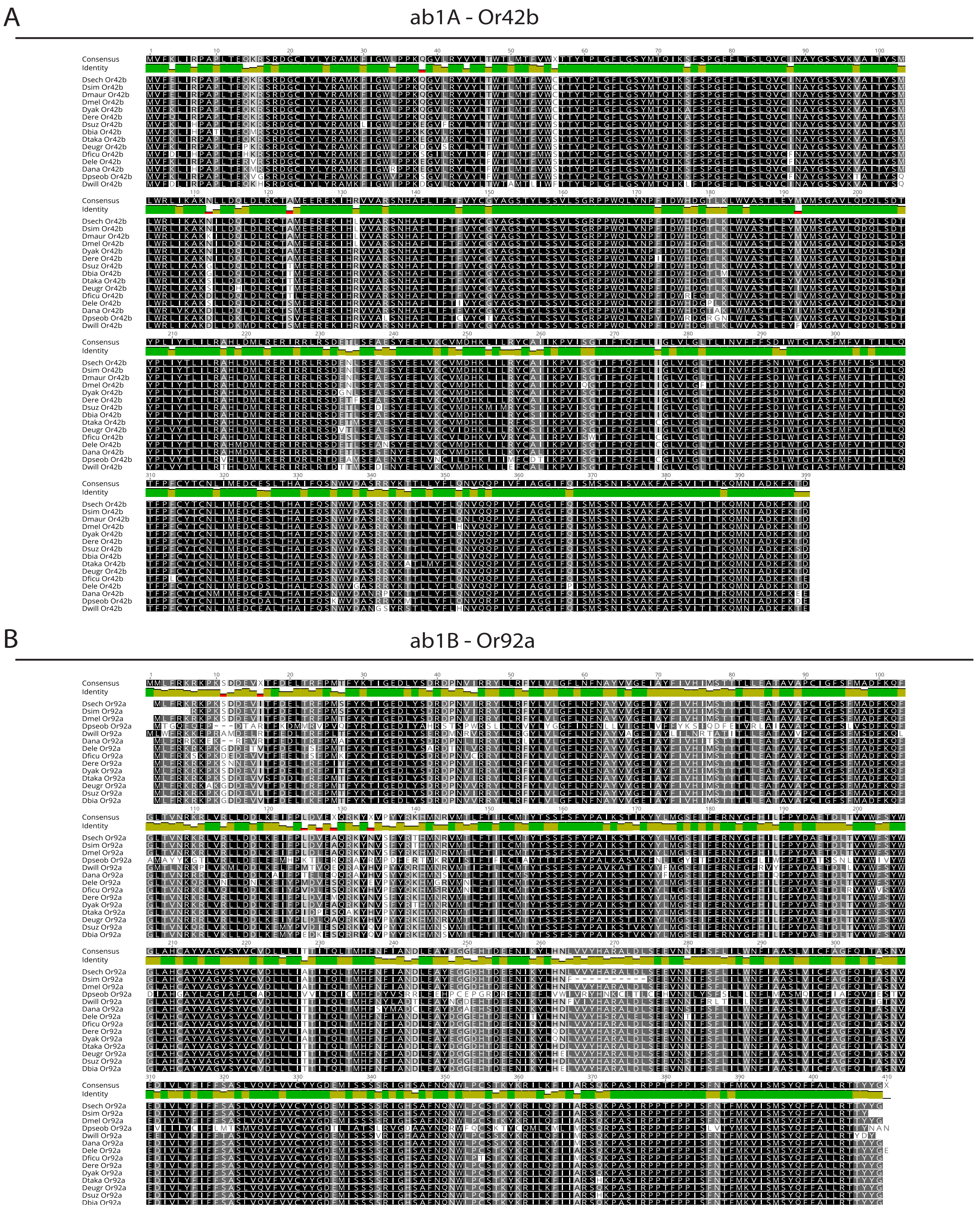


**Supplementary Figure 3. Protein sequence alignments for ab1 across *Sophophora*.**
Two protein alignments for those that are housed in the ab1 sensillum type (Or42b and Or92a) are displayed for as many species as were available for comparison. Highly similar amino acids across species are displayed in black, while dark grey, light grey and white positions denote increasing variability in sequence data. Both of these receptors display high functional conservation of olfactory function across all 20 species for which SSR data was generated, and maintain identical odorant profiles to those described from *D. melanogaster* adults.

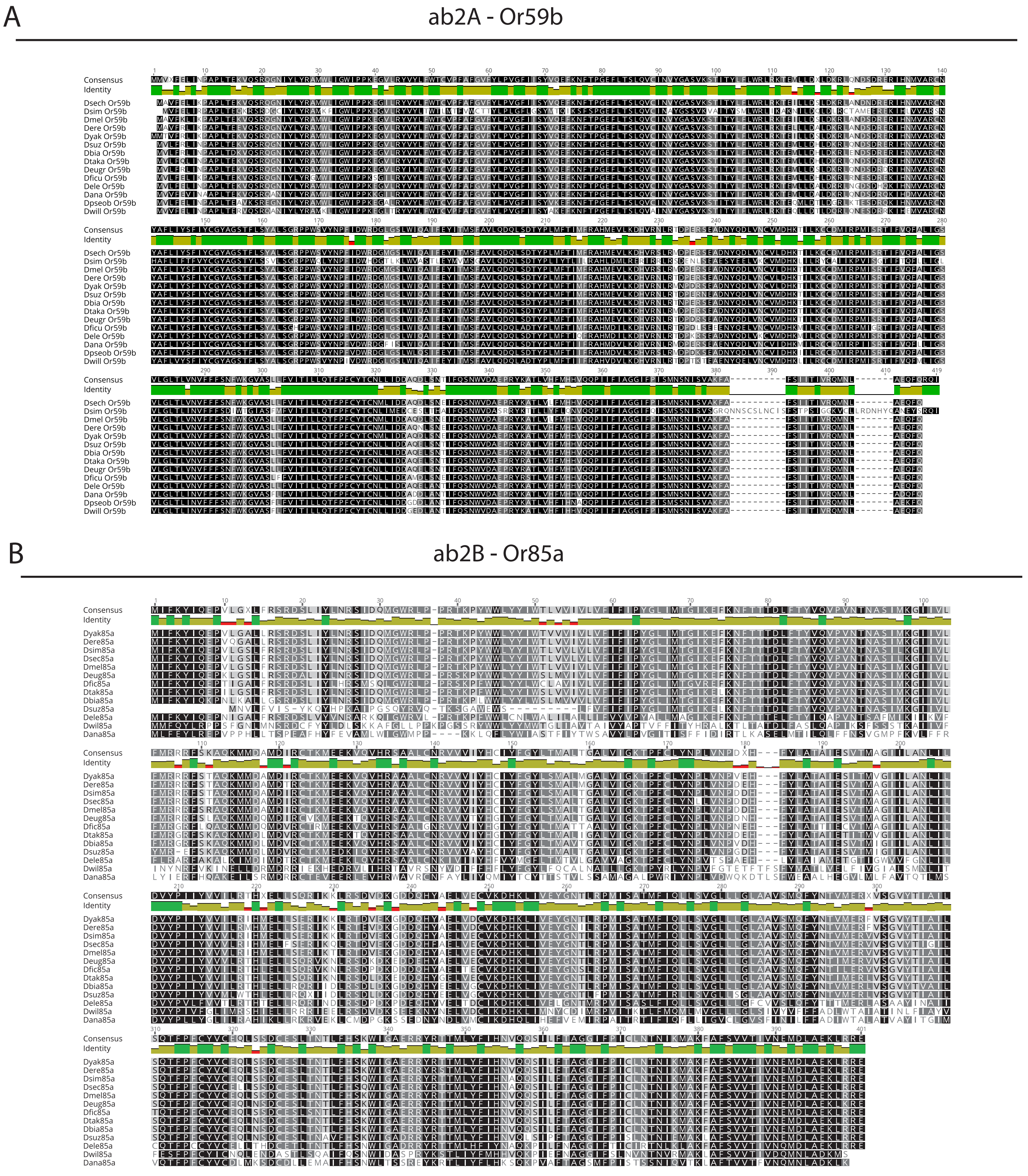


**Supplementary Figure 4. Protein sequence alignments for ab2 across *Sophophora*.**
Two protein alignments for those that are housed in the ab2 sensillum type (Or59b and Or85a) are displayed for as many species as were available for comparison. Highly similar amino acids across species are displayed in black, while dark grey, light grey and white positions denote increasing variability in sequence data. Only one of these receptors displays high functional conservation of olfactory function across all 20 species for which SSR data was generated (Or59b), and maintains identical odorant profiles to those described from *D. melanogaster* adults. However, the other (Or85a) produced highly variable olfactory response profiles in SSR data for 7 of the 20 examined species, including *D. willistoni*, *D. affinis*, *D. pseudoobscura*, *D. ananassae*, *D. birchii*, *D. elegans*, and *D. suzukii* adults.

A

ab3A - Or22a

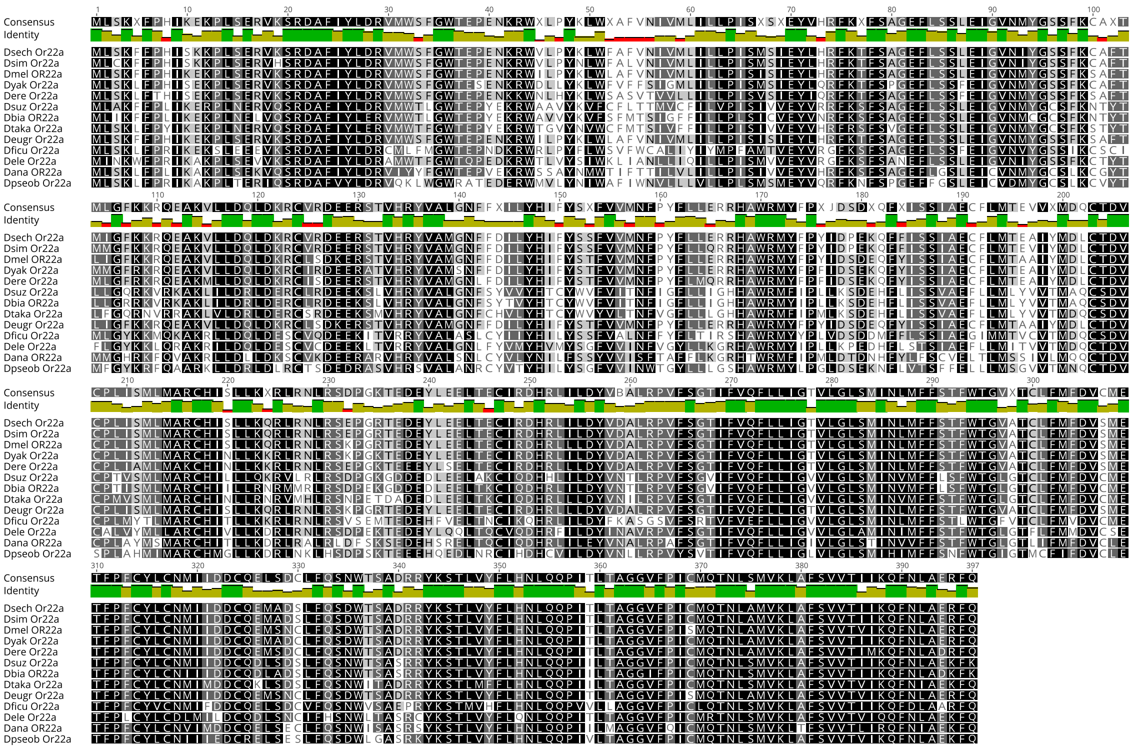

B

ab3B - Or85b

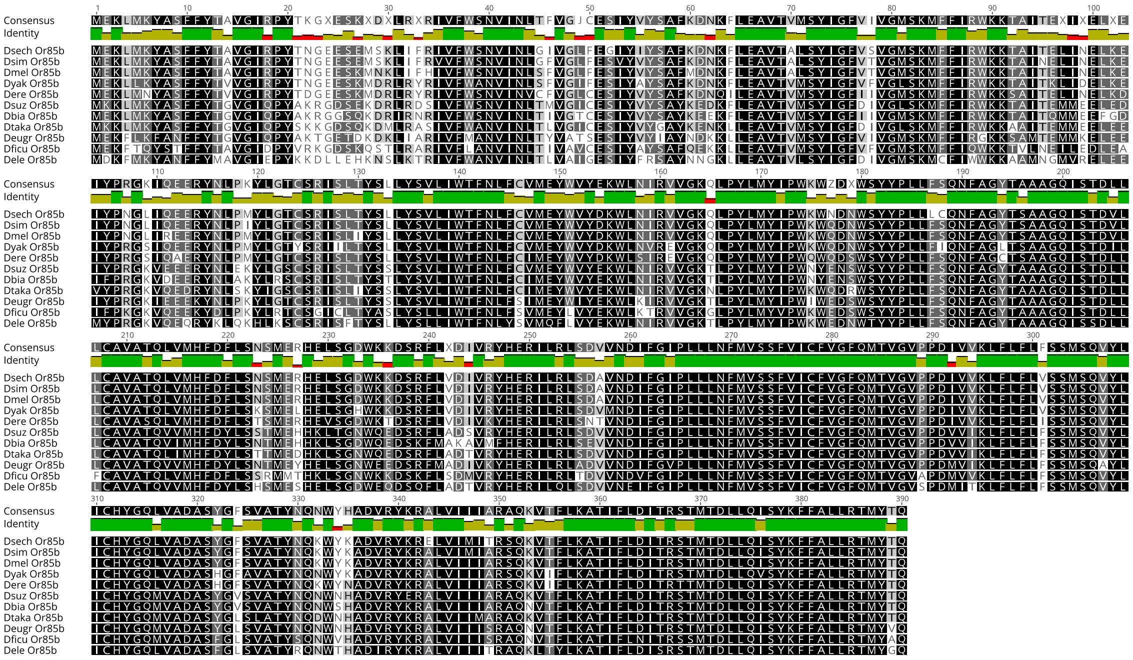

**Supplementary Figure 5. Protein sequence alignments for ab3 across *Sophophora*.**
Two protein alignments for those that are housed in the ab3 sensillum type (Or22a and Or85b) are displayed for as many species as were available for comparison. Highly similar amino acids across species are displayed in black, while dark grey, light grey and white positions denote increasing variability in sequence data. Only one of these receptors displays high functional conservation of olfactory function across all 20 species for which SSR data was generated (Or85b), and maintains identical odorant profiles to those described from *D. melanogaster* adults. However, the other (Or22a) produced highly variable olfactory response profiles in SSR data for 13 of the 20 examined species, at least in comparison to the *D. melanogaster* model, including *D. willistoni*, *D. affinis*, *D. subobscura*, *D. pseudoobscura*, *D. elegans*, *D. ficusphila*, *D. eugracilis*, *D. pseudotakahashii*, *D. biarmipes*, *D. subpulchrella*, *D. suzukii*, *D. simulans* and *D. sechellia* adults.

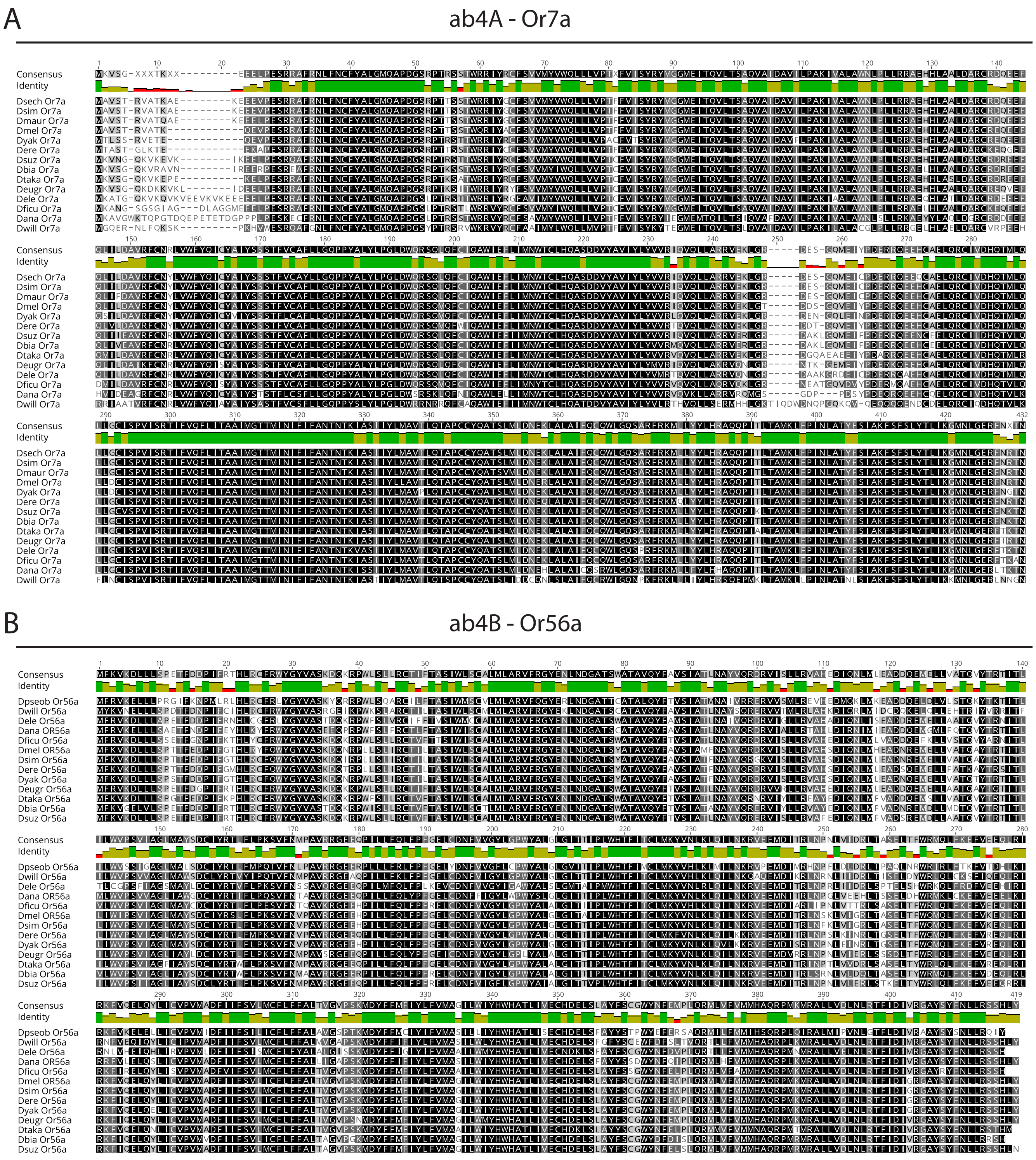


**Supplementary Figure 6. Protein sequence alignments for ab4 across *Sophophora*.**
Two protein alignments for those that are housed in the ab4 sensillum type (Or7a and Or56a) are displayed for as many species as were available for comparison. Highly similar amino acids across species are displayed in black, while dark grey, light grey and white positions denote increasing variability in sequence data. Both of these receptors display high functional conservation of olfactory function across all 20 species for which SSR data was generated, and maintain identical odorant profiles to those described from *D. melanogaster* adults.

tree of Or22a (whole length) similarity

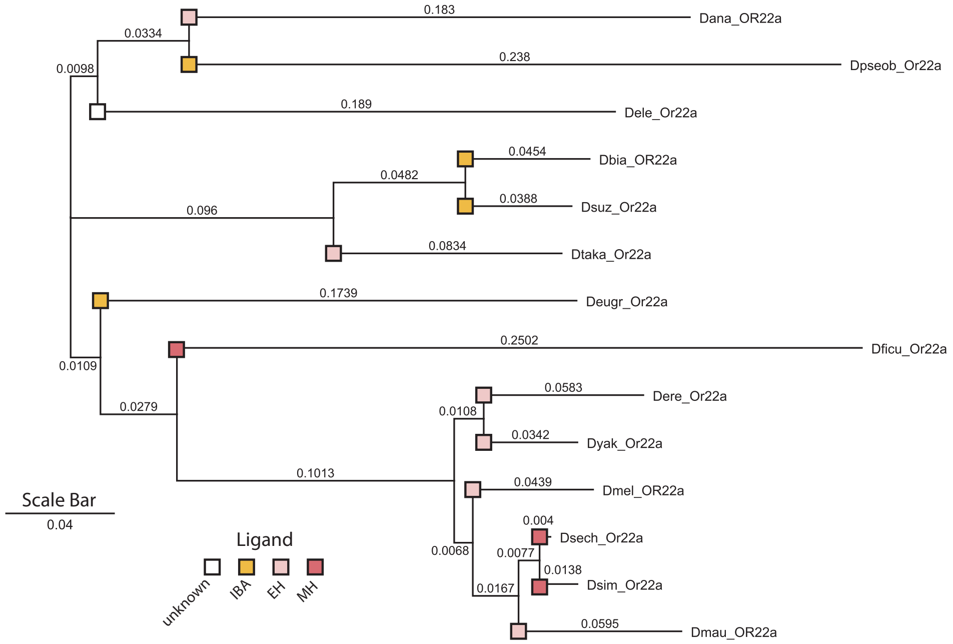


tree of Or85a (whole length) similarity

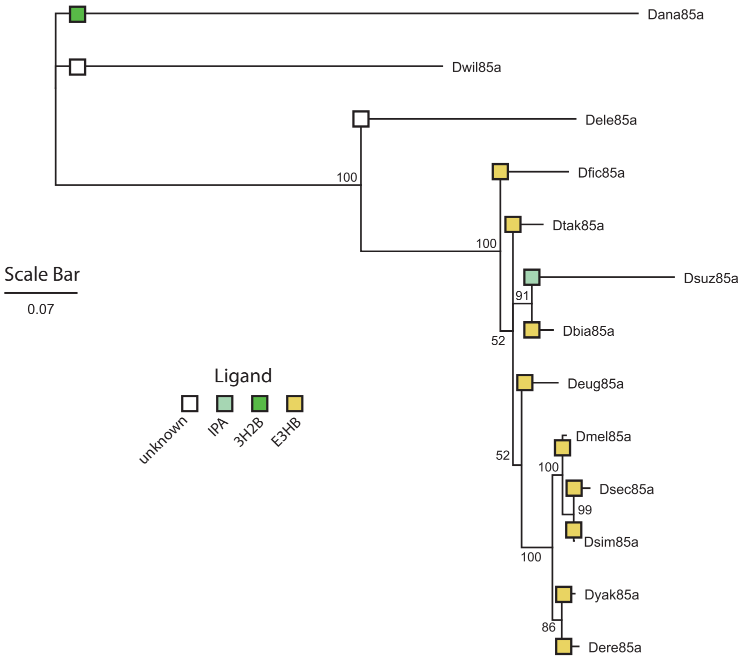


**Supplementary Figure 7. Trees of Or22a & Or85a sequence similarity across available species.**
Displayed is the whole length trees of similarity for both Or22a and Or85a for each of the Sophophora species for which genomic data was available. Also shown are the functional receptor ligands for each species, including isobutyl acetate (IBA), ethyl hexanoate (EH) and methyl hexanoate (MH) for Or22a, as well as isopentyl acetate (IPA), 3-hydroxy-2-butanone (3H2B; acetoin), and ethyl-3-hydroxybutanoate (E3HB). It appears that receptor ligands are shared across many species, and mirror phylogenetic relationships for this subgenus. Moreover, it would appear that ligand shifts have occurred across multiple subgroups, including the presence of EH in 3 different branches (i.e. *D. ananassae*, *D. takahashii*, as well as the *melanogaster* clade) for Or22a. Similarly, the evolution of IBA has also arisen in several subgroups (including: *D. pseudoobscura*, *D. eugracilis*, and the *suzukii* clade).

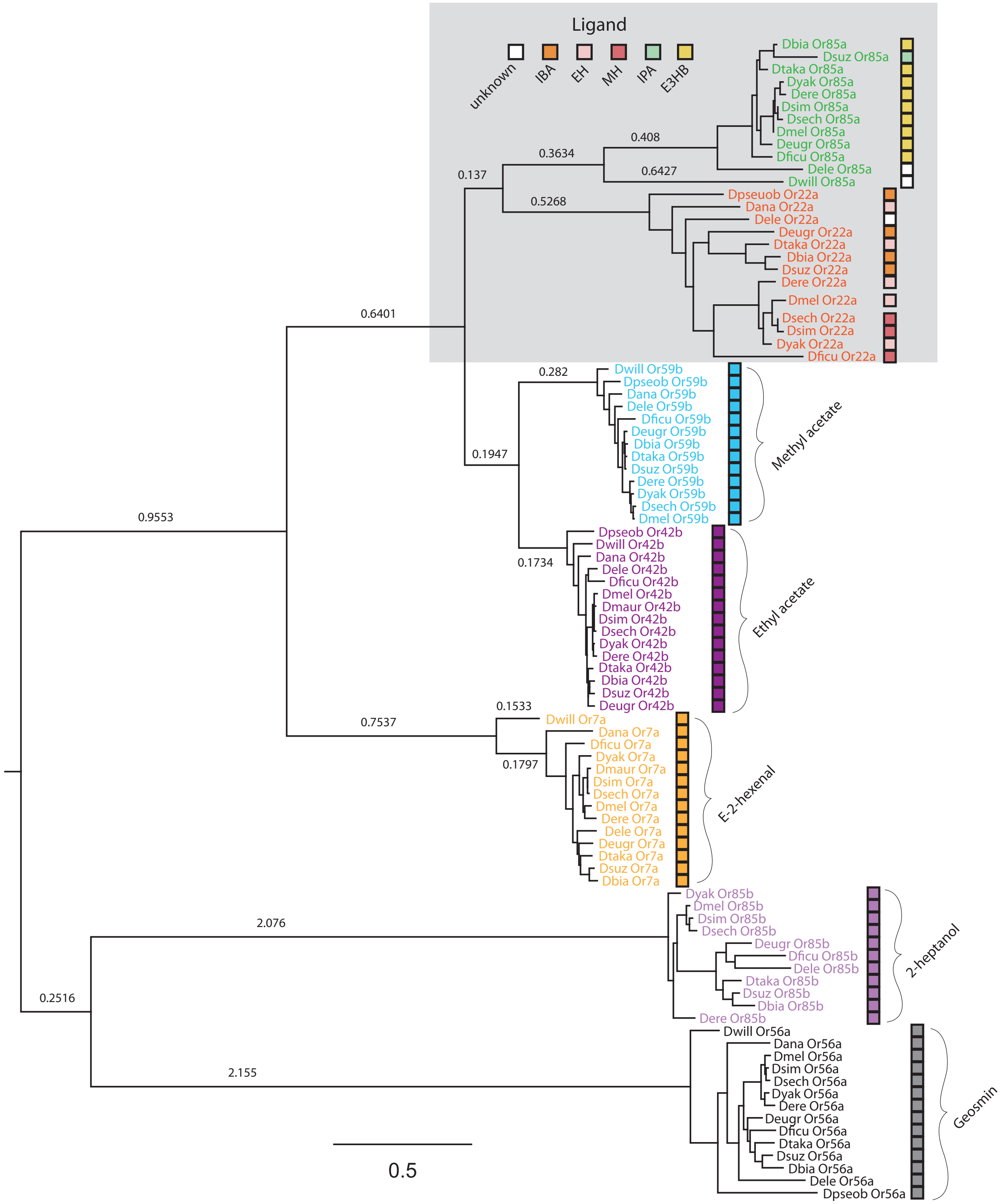


### Supplementary Figure 8. Tree of all receptor sequence orthologues

Shown is the tree of sequence similarity for all receptors addressed in this study. Five sets of sequences all provided identical single sensillum recordings (SSR) in regards to best ligand, and we also noted very little change in sequence data for these olfactory receptors (e.g. Or59b, Or42b, Or7a, Or85b, and Or56a). However, two sets of sequences were more variable, including our Or22a and Or85a data. Here we also noted a strong variation in both ligand and olfactory tuning for these receptors when compared across species, which is further reflected in the increased distances between species for the same receptor.

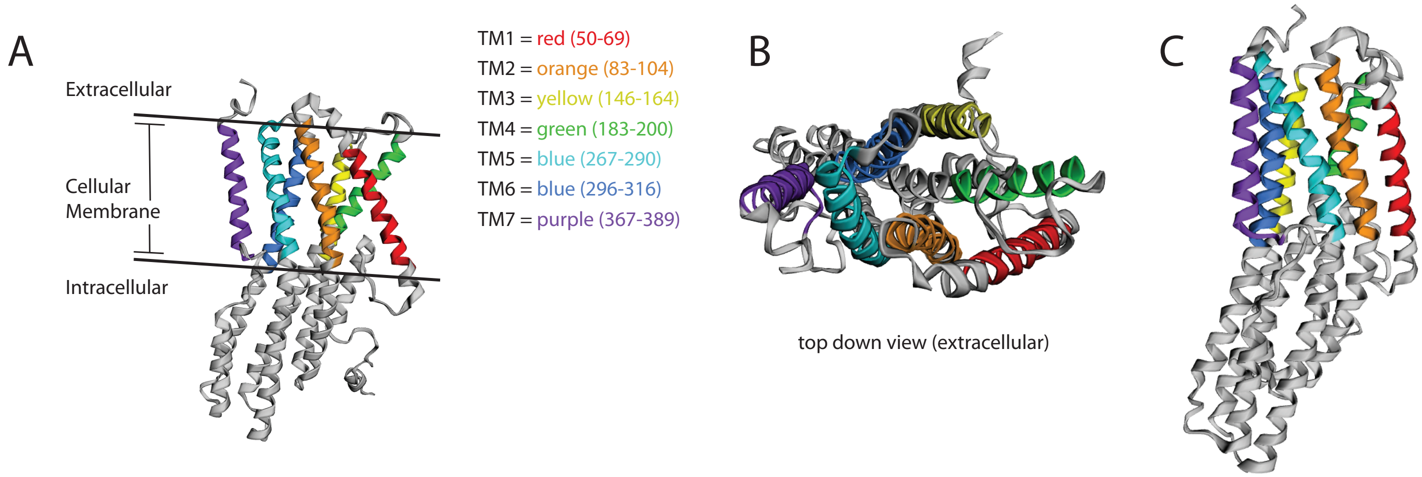


**Supplementary Figure 9. Protein tertiary structure for Or22a from *D. melanogaster*.**
**(A)** Depicted is the complete protein structure of Or22a, with each of the seven transmembrane domains (TM1-7) shown in color code. **(B)** Top-down view of the Or22a tertiary protein structure, highlighting the overlap of TM1 and TM4, where the putative binding pocket is located for this olfactory receptor [14]. **(C)** Rotated view again highlighting the transmembrane domains, as well as the 3-dimensional orientation of each domain.

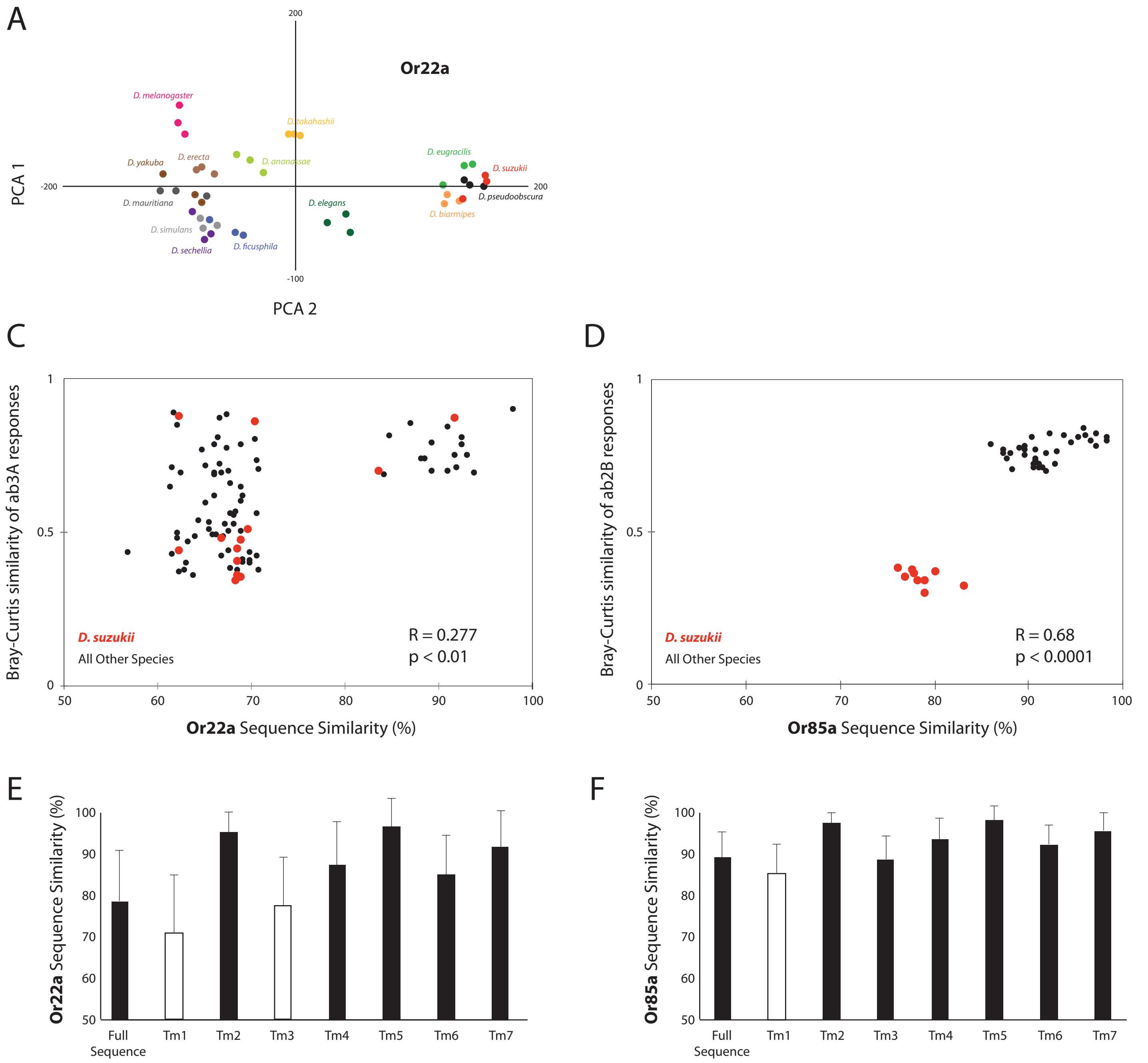


**Supplementary Figure 10. SSR-responses and their correlation with receptor sequence.**

(A-B) Principal component analysis (PCA) based on SSR responses from the ab3A- and ab2B-like sensilla of 14 Drosophilid species for which sequence data was available. (C) Correlations between pairwise similarities of Or22a sequences and the corresponding pairwise similarities of ab3A sensillum SSR responses. Here *D. suzukii* does not cluster outside the others, suggesting that a similar Or22a is common to all examined species. (D) Correlations between pairwise similarities of Or85a sequences and the corresponding pairwise similarities of ab2B sensillum responses (R- and P-values calculated by Mantel test for comparison of matrices). Here *D. suzukii* does in fact cluster outside all the other species, suggesting a large deviation in both SSR and sequence data. (E-F) Average pairwise sequence similarities within the Or22a homologs (E) and the Or85a homologs (F) as well as the corresponding seven transmembrane regions (Tm1-7) of both receptor proteins. Here we observe that Tm1 and Tm3 appear critical for Or22a functional variation, while just Tm1 for Or85a data appears to vary, though we continue to suggest that Or85a has been replaced in *D. suzukii* and other highly variable species (Figure 4 C).

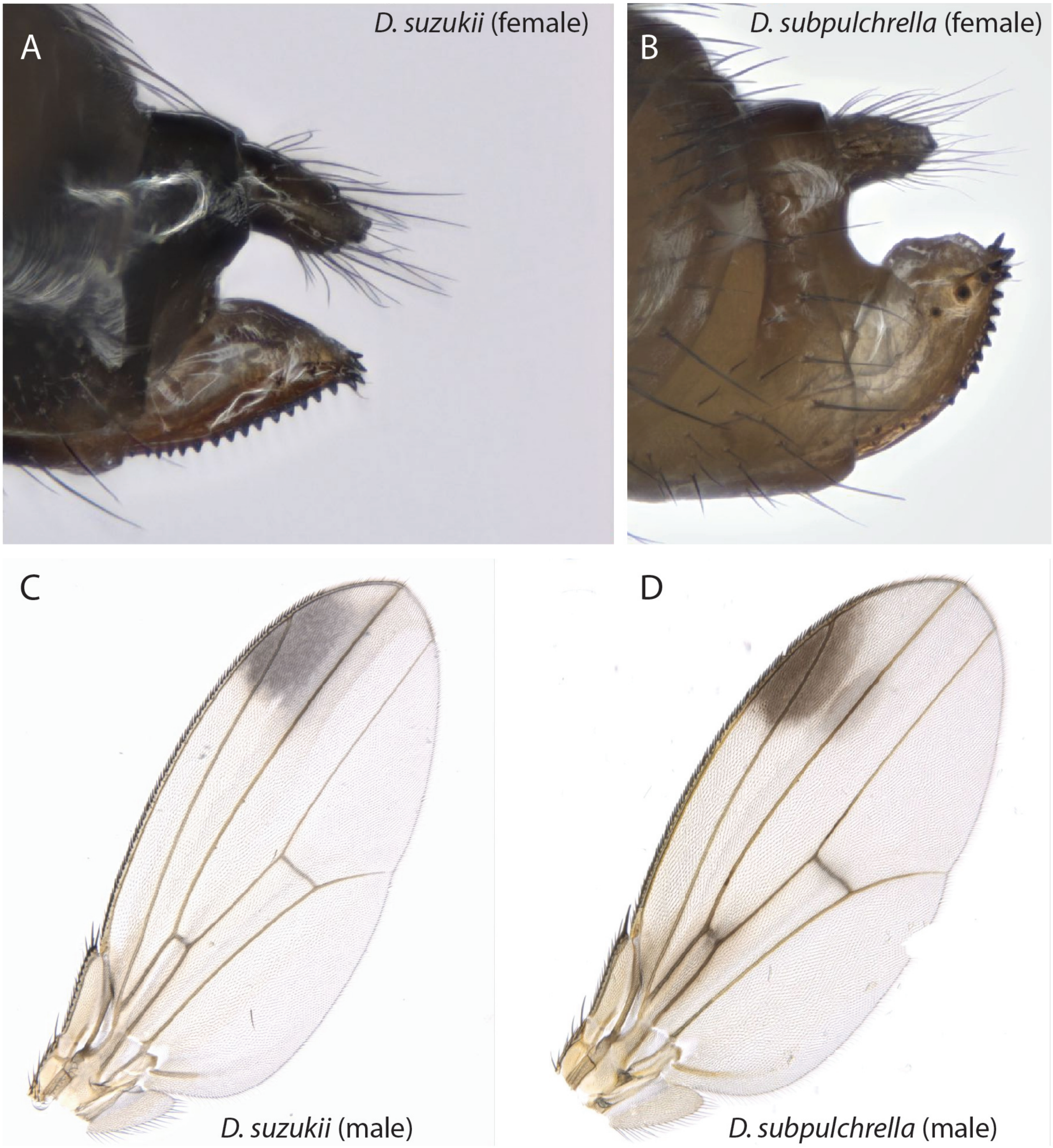


**Supplementary Figure 11. Comparison of *D. suzukii* with *D. subpulchrella*, the closest relative.**
**(A)** Lateral view of the female ovipositor of *D. suzukii* adult. Note the high number of heavily sclerotized teeth, and overall the heavy level of darkened scleritization of the ovipositor. **(B)** Lateral view of the female ovipositor of D. *subpulchrella* adult. Note the reduced number and size of teeth. Also shown are the wings of male *D. suzukii* adults **(C)** as well as those from male *D. subpulchrella* adults **(D)**, where the latter contains a more proximal pigmentation, and two slightly separated wing spots.

A

ab1C - Gr21a

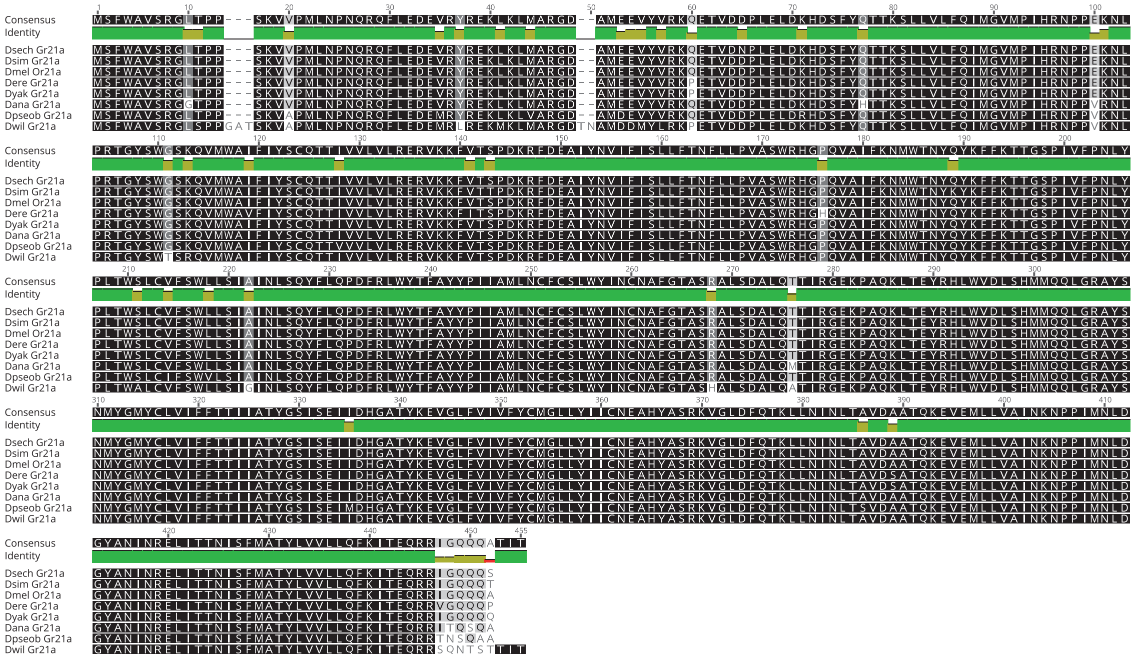

B

ab1C- Gr63a

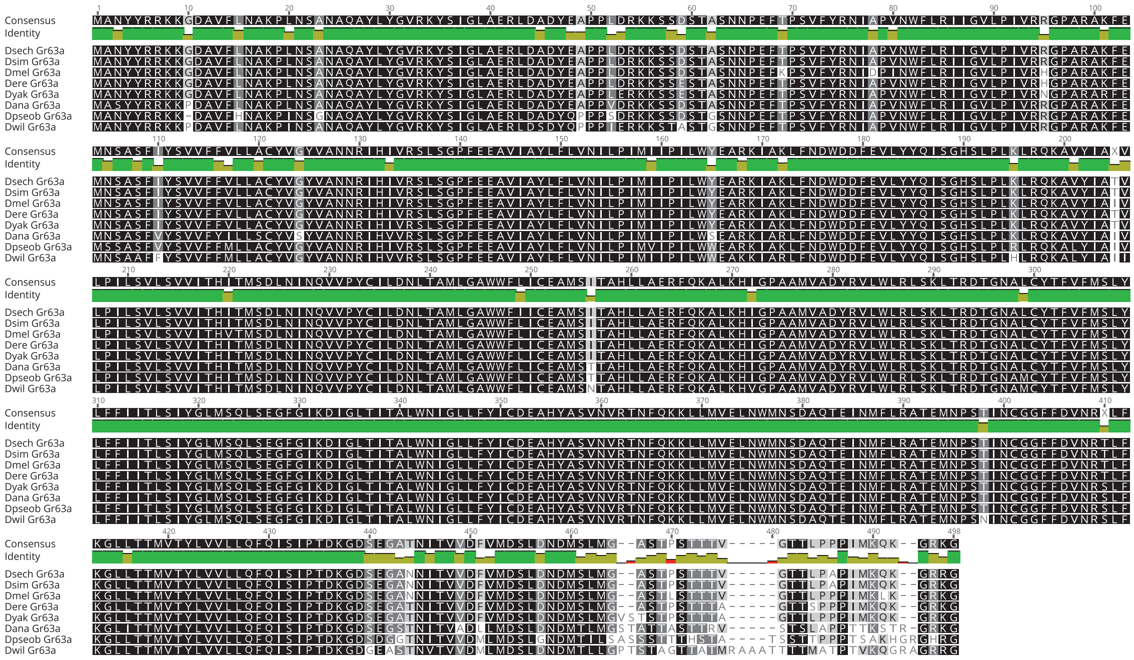

**Supplementary Figure 12. Protein sequence alignments for ab1C across *Sophophora*.**
Two protein othologue alignments for those gustatory receptors that are housed in the ab1 sensillum type (Gr21a and Gr63a) are shown for as many species as were available for comparison (see Robertson et al. 2009; doi: 10.1673/031.009.1901). Highly similar amino acids across species are displayed in black, while dark grey, light grey and white positions denote increasing variability in amino acid data. Both of these receptors display high conservation of olfactory function across all 20 species for which SSR data was generated, and maintain identical odorant profiles to those described from *D. melanogaster* adults; however, species do vary in their sensitivity to carbon dioxide (the best ligand).
